# Multimodal Gradient Mapping of Rodent Hippocampus

**DOI:** 10.1101/2021.11.22.469439

**Authors:** Brynja Gunnarsdóttir, Valerio Zerbi, Clare Kelly

## Abstract

The hippocampus plays a central role in supporting our coherent and enduring sense of self and our place in the world. Understanding its functional organisation is central to understanding this complex role. Previous studies suggest function varies along a long hippocampal axis, but there is disagreement about the presence of sharp discontinuities or gradual change along that axis. Other open questions relate to the underlying drivers of this variation and the conservation of organisational principles across species. Here, we delineate the primary organisational principles underlying patterns of hippocampal functional connectivity (FC) in the mouse using gradient analysis on resting state fMRI data. We further applied gradient analysis to mouse gene co-expression data to examine the relationship between variation in genomic anatomy and functional organisation. Two principal FC gradients along a hippocampal axis were revealed. The principal gradient exhibited a sharp discontinuity that divided the hippocampus into dorsal and ventral compartments. The second, more continuous, gradient followed the long axis of the ventral compartment. Dorsal regions were more strongly connected to areas involved in spatial navigation while ventral regions were more strongly connected to areas involved in emotion, recapitulating patterns seen in humans. In contrast, gene co-expression gradients showed a more segregated and discrete organisation. Our findings suggest that hippocampal functional organisation exhibits both sharp and gradual transitions and that hippocampal genomic anatomy exerts a subtle influence on this organisation.

## 1. Introduction

As the seat of learning and memory, and a nexus for the integration of time, space, and emotion, the hippocampus is crucial to our coherent and enduring sense of self and our place in the world (Chersi and Burgess, 2015; Fink, 2019; Phelps, 2004; Strange et al., 2014). A better appreciation of the functional organisation of the hippocampus, in particular, of its connectivity with the rest of the brain, is needed to advance our understanding of its pivotal role in these diverse cognitive processes. Moreover, such an understanding is fundamental to discovering how hippocampal dysfunction gives rise to the devastating disruptions of mnemonic, emotional, and cognitive processing that characterize neurological and psychiatric conditions such as Alzheimer’s disease, depression, and schizophrenia (Andersen et al., 2007).

The hippocampus is a phylogenetically preserved structure with extensive connections throughout the brain. Its main inputs and outputs are via the entorhinal cortex and the fornix, but it also projects directly to the amygdala and receives direct projections from the locus coeruleus. Through the entorhinal cortex, the hippocampus connects with cortical regions such as the peri-, and postrhinal cortices, olfactory structures, piriform, agranular insular, medial prefrontal, anterior cingulate and retrosplenial cortex. The entorhinal cortex additionally receives input from subcortical structures such as the amygdala and projects to the striatum, especially the nucleus accumbens, via the subiculum. The fornix pathway links the hippocampus to basal forebrain, hypothalamic and brainstem regions. Through this pathway the hippocampus connects with septal nuclei and the mamillary bodies of the hypothalamus and projects, via the subiculum, to anterior thalamic nuclei and the ventral striatum (Andersen et al., 2007; Wagatsuma et al., 2018).

How do these diverse anatomical connections enable the complex functions of the hippocampus? Two primary principles are proposed to govern its functional organisation. The first follows the transverse axis of the hippocampus along the proximal-distal axis of its subfields and is based on cytoarchitectural distinctions. This organisational principle divides the hippocampus into its subfields (DG, CA3, CA2 and CA1) and reflects the unique excitatory flow of information through the hippocampal formation - the trisynaptic circuit (Andersen et al., 2007). The second organisational principle proposes that hippocampal function varies along its long axis, which runs anterior-to-posterior in humans and ventral-to-dorsal in rodents (see **Figure 1** for a schematic overview of the mouse hippocampus).

**Figure 1:**
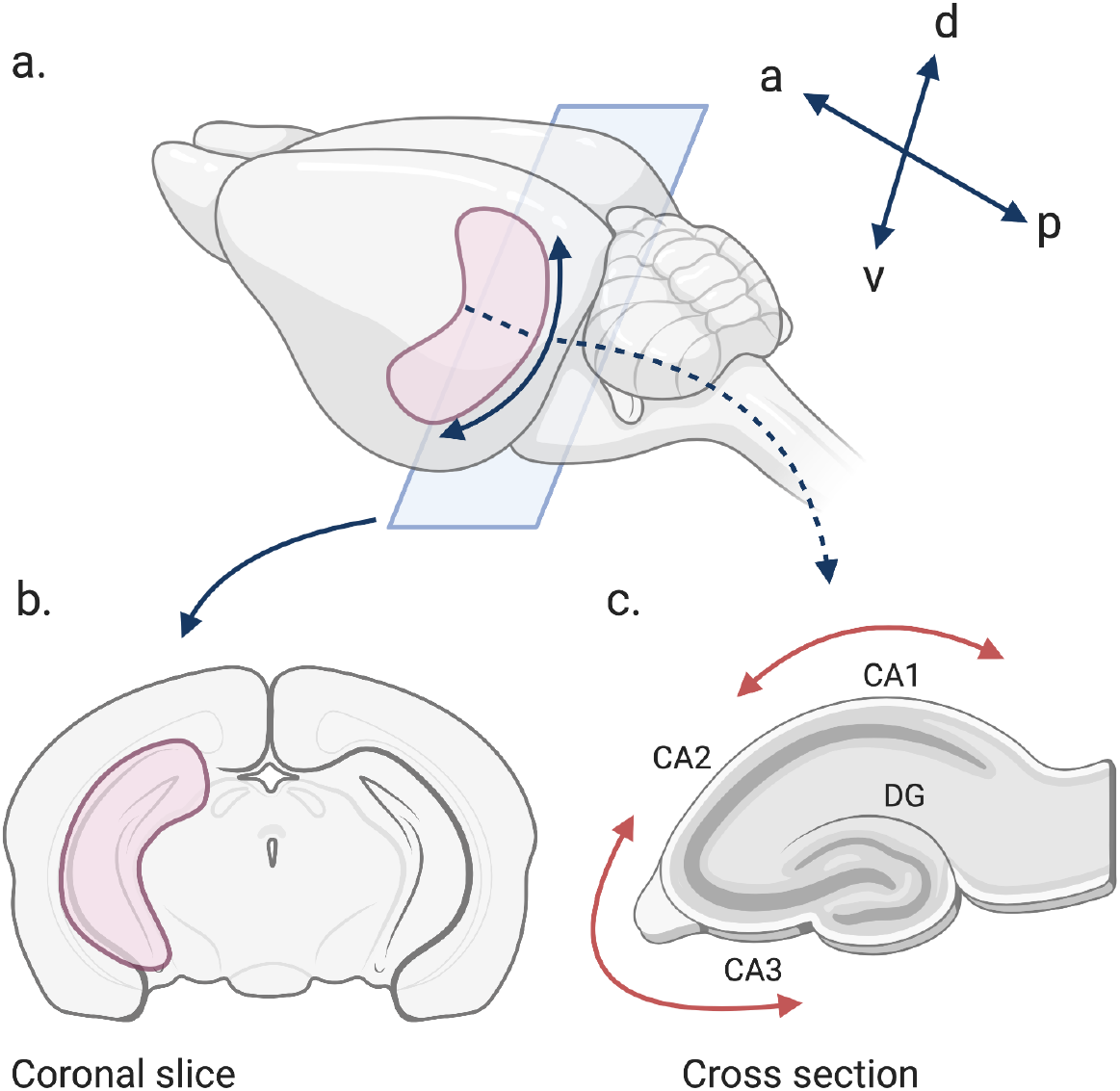
Schematic of Mouse Hippocampus. a. Shows the hippocampus in the mouse brain in red, extending in a C-shape along a long (dorso-ventral; d-v) axis, from the midline of the brain to the incipient temporal lobe(a and p denote the anterior-to-posterior axis). b. Shows a coronal slice of the mouse brain where the hippocampus is labelled red. c. Shows a cross section of the hippocampus where the dentate gyrus (DG), and cornu ammonis (CA) subfields are labelled. Red arrows denote the proximal-to-distal axis, followed clockwise here, of the hippocampal subfields.

A traditional view has been that function along the long axis of the hippocampus is clearly segregated into dorsal and ventral compartments, such that the ventral hippocampus is associated with emotion and fear, based on denser reciprocal connectivity with the amygdala and hypothalamus (Strange et al., 2014), while the dorsal hippocampus is associated with spatial navigation and memory, based partly on stronger input from the sensory cortices (Moser and Moser, 1998). A recent fMRI-based parcellation of the subcortex divided the hippocampus into anterior and posterior compartments at finer scales of parcellation (Tian et al., 2020). Gene expression (GE) studies in rodents similarly characterise the genomic anatomy of the hippocampus as parcellated, such that the hippocampal subfields are subdivided along the long axis into distinct molecular domains (Bienkowski et al., 2018; Dong et al., 2009; Fanselow and Dong, 2010; Floriou-Servou et al., 2018; Thompson et al., 2008). Complicating this picture, the subfields appear to differ in genomic anatomy, so that the difference in GE along the long axis of CA1 is smaller or more graded compared to CA3, which is composed into discrete molecular domains (Cembrowski et al., 2016; Dong et al., 2009).

Anatomical tract tracing studies examining structural connectivity in rodents provide further support for gradual organisational variation along the long axis. These studies suggest a topographical gradient such that projections from the hippocampus to cortical and subcortical structures, such as the cingulate cortex, nucleus accumbens, and amygdala, change gradually along the hippocampal long axis (Kishi et al., 2006; Strange et al., 2014). Even more recent studies suggest that the boundary-or-gradient question may be misconceived and that a nuanced picture of hippocampal organisation is needed. For example, a recent rodent study combining GE and anatomical tracing found that while discrete molecular domains of the hippocampus were connected to independent brain-wide networks, there was also evidence of distributed hippocampal contributions, with the latter likely supporting the integration of information across distinct hippocampal networks (Bienkowski et al., 2018).

Here, we applied gradient analysis to shed further light on the complex organisation of the hippocampus. Gradient analysis characterises the main axes of variance in numerical data in a location-dependent pattern; in doing so, it can capture both sharp and gradual changes within a structure and can tease out overlapping principles of organisation (Przeździk et al., 2019). Previous studies have found that gradient analysis captures known variation in microstructure, structural connectivity, GE maps, and task based fMRI results, and can extend findings from previous studies (Fulcher et al., 2019; Haak et al., 2018; Huntenburg et al., 2018; Margulies et al., 2016; Marquand et al., 2017; Paquola et al., 2019).

In humans, gradient analyses of the hippocampus have uncovered a principal gradient of organisation such that FC changes in a gradual manner along the hippocampus long axis (Vos de Wael et al., 2018; Przeździk et al., 2019). These studies reported stronger connectivity of the anterior hippocampus to regions associated with higher-order cognition and emotion (e.g., default network) and of posterior regions to visual and sensory networks regions. Vos de Wael et al. (2018) additionally uncovered a second medial-to-lateral gradient, which followed hippocampal infolding, and was correlated with surface-based markers of intracortical microstructure, indicating interactions between tissue microstructure and function.

We set out to further advance our understanding of hippocampal functional organisation using the gradient analysis framework proposed by Haak et al. (2018) to map the main axes of variation in patterns of functional connectivity between the hippocampus and the rest of the brain, based on rsfMRI data collected from lightly anaesthetized mice. Our analysis applied precisely the same methods to the mouse data as are used for human data, thus facilitating direct comparison between principles of functional organisation across species. Capitalising on this translational approach, we applied the same gradient analysis to hippocampal gene coexpression (GCE) data obtained using the Anatomic Gene Expression Atlas (AGEA) to compare the main organisational principles of hippocampal FC to patterns of GCE (Ng et al., 2009). Previous studies examining differentially expressed genes within the hippocampus have found strongly represented functional categories that could be important for the establishment and maintenance of FC (Thompson et al., 2008). Bienkowski et al. (2018) found distinct anatomical connectivity for distinct hippocampal GE domains in rodents. Additionally, a recent study by Vogel et al. (2020) found a strong relationship (r^2^=0.40) between a human hippocampal FC gradient and the predicted location of hippocampal tissue samples based on GE patterns. We hypothesized that these analyses would uncover a gradual long axis FC gradient of mouse hippocampal FC, analogous to the gradients described in humans (Vos de Wael et al., 2018; Przeździk et al., 2019). We expected to find corresponding long axis variation in GCE as well, demonstrating a role for genomic anatomy in setting FC topography.

## 2. Methods and Materials

### 2.1 Animals, rsfMRI Preparation

A pre-existing dataset comprising rsfMRI acquired scans from 50 lightly anesthetized C57Bl6/J and wild type mice was used in this study (Filipello et al., 2018; Zerbi et al., 2015). All mice were adults (age: 14+/-4 weeks; bodyweight: 25.1+/-3 g; 39/11 male/female). The experiments were performed in accordance with the Swiss federal guidelines for the use of animals in research, and under licensing from the Zürich Cantonal veterinary office.

The mice were prepared for rsfMRI using established protocols (Chelini et al., 2019; Grandjean et al., 2014; Zerbi et al., 2015). Anaesthesia was induced with 4% isoflurane and the animals were endotracheally intubated and the tail vein cannulated. The mice were positioned on an MRI-compatible cradle, and artificially ventilated at 80 breaths per minute with a 1:4 O_2._ to air ratio, and a 1.8 ml/h flow (CWE, Ardmore, USA). A bolus injection of pancuronium bromide (0.2 mg/kg), a muscle relaxant, was administered, and isoflurane was reduced to 1%. Throughout the experiment, the mice received a continuous infusion of pancuronium bromide (0.4 mg/kg/h). Body temperature was monitored using a rectal thermometer probe, and maintained at 36.5 ± 0.5 °C.

### 2.2 rsfMRI

Data was acquired using a Biospec 70/16 small animal MR system (Bruker Biospin MRI, Ettlingen, Germany) equipped with a cryogenic quadrature surface coil for signal detection (Bruker BioSpin AG, Fällanden, Switzerland). Standard adjustments included the calibration of the reference frequency power and the shim gradients using MapShim (Paravision v6.1). A standard gradient-echo echo planar imaging sequence (GE-EPI, repetition time *TR* = 1 s, echo time *TE* = 15 ms, in-plane resolution *RES* = 0.22 × 0.2 mm^2^, number of slice *NS* = 20, slice thickness *ST* = 0.4 mm, slice gap = 0.1 mm) was used to acquire 900 volumes in 15 minutes.

### 2.3 Preprocessing

Resting state fMRI (rsfMRI) datasets were preprocessed with an existing automated pipeline, adapted for the mouse (Zerbi et al., 2015). This pipeline includes independent component analysis (ICA) based artefact removal (FSL’s FIX https://fsl.fmrib.ox.ac.uk/fsl/fslwiki/FIX), band-pass filtering (0.01-0.25 Hz), skull-stripping and normalization to the Allen Mouse Brain Common Coordinate Framework (CCFv3) (Wang et al., 2020). Normalization was performed through linear affine and nonlinear greedy SyN diffeomorphic transformation metric mapping using ANTs v2.1 (picsl.upenn.edu/ANTS). Normalized, preprocessed volumes were downsampled to match the native resting state EPI resolution.

### 2.4 Gradient Analysis

We used the data-driven connectopic mapping method (Haak et al., 2018) to examine FC gradients in the mouse hippocampus. This method is suitable for analysing both volume (permitting the inclusion of the subcortex) and surface (cortex) data and has previously been used to examine the organization of the human hippocampus (Przeździk et al., 2019).

Connectopic mapping extracts the principal modes of FC change between a predefined region of interest (ROI) and the rest of the brain. First, connectivity “fingerprints” are computed, quantifying FC between each voxel in the region of interest (ROI, here, the hippocampus) and every other voxel in the brain using Pearson correlation. Next, a similarity matrix is computed which describes the pairwise similarities between the connectivity fingerprints of all the voxels within the ROI, using the η^2^ coefficient. Finally, manifold learning, using the Laplacian Eigenmaps (LE) algorithm, finds a low-dimensional representation of the similarity matrix. The low-dimensional representations are eigenvectors, referred to as FC gradients. This analysis is performed at the level of individual participants; group analyses are performed by averaging individual similarity matrices across subjects into a group similarity matrix, then applying the manifold learning to that group-level matrix (Haak et al., 2018).

The gradients represent the dominant modes of change in whole-brain FC patterns across the ROI, whereby similar values within a gradient signify similar connectivity patterns (Haak et al., 2018). Because they correspond to eigenvectors, the gradients are sorted such that the first gradient captures the highest amount of information on FC variation, the second the second-highest and so on (Navarro Schröder et al., 2015).

Separate right and left hippocampus masks were created using the CA1, CA2, CA3 and DG ROIs as defined in the Allen Common Coordinate Framework (CCF) (V3, http://help.brain-map.org/download/attachments/2818169/MouseCCF.pdf). A whole-brain mask was created by including all CCF ROIs that were present in all mice (i.e., areas of signal loss/dropout were excluded) (see **Supp. Figure 1**). First, we computed FC between each hippocampal voxel (left *n* = 441; right *n* = 436) and all other voxels within the whole-brain mask for each mouse. Next, similarity matrices were computed, averaging across individuals to create a group similarity matrix, and gradient analysis was conducted using scripts available at https://github.com/koenhaak/congrads. Analyses were performed for the left and right hippocampus separately. We chose to investigate the three principal gradients for each hemisphere further, based on the clearest elbow in Scree Plots derived from a group gradient analysis on an exploration dataset (*n* = 30) (See **Supp. Figure 2** and **3** and **S1** in the **Supplementary Material** for further information on the Scree Plot method).

We assessed the consistency and stability of the gradients by computing the spatial correlation between the first three gradients across exploration (*n* = 30) and validation (*n* = 20) datasets. The first FC gradient revealed a sharp change in connectivity along the hippocampal long axis, suggesting functional segregation. To examine whether this reflected a functionally distinct region dorsally, we constructed a graph showing gradient value as a function of distance from the most dorso-medial location of the hippocampus (following Przeździk et al. (2019). All analyses were conducted using Spyder through the Anaconda distribution or fsl in the Python programming language (*Anaconda Software Distribution*, 2020; Rossum and Jr, 2011; Woolrich et al., 2009).

### 2.5 Bootstrap Aggregation of Clusters and Consensus Clustering

To visualise the patterns of FC contributing to the gradients detected, the FC gradients were clustered, and seed-based FC was computed for the resultant clusters (see **Figure 2** for a schematic of the methods). For comparison with patterns of FC obtained using traditional anatomical divisions, FC analysis was also conducted using the primary hippocampal anatomical compartments (CA1, CA2, CA3, and DG) as seeds.

**Figure 2:**
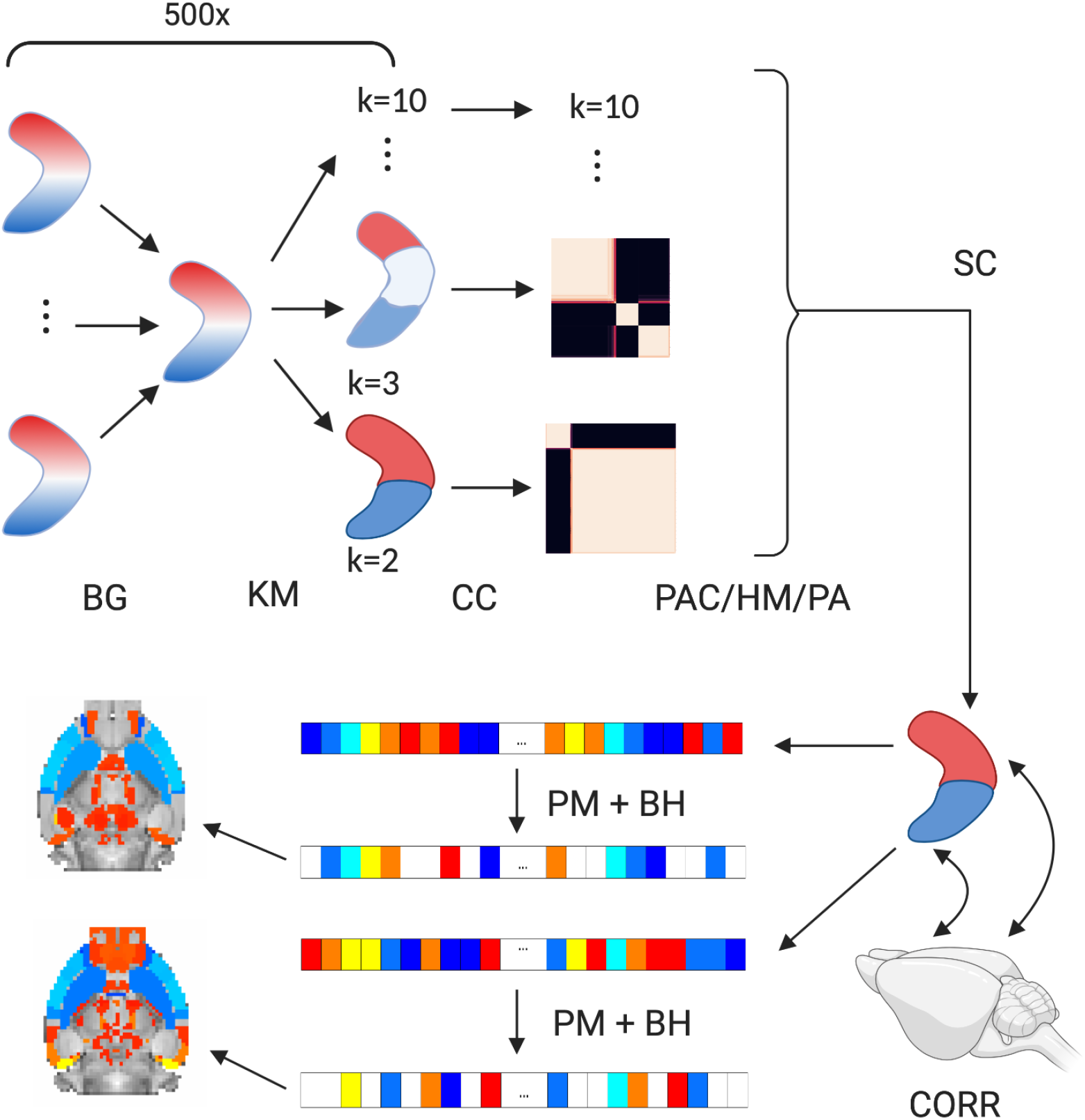
Schematic Overview of Consensus Clustering and Seed-based FC analysis. Gradients were resampled with replacement by choosing random gradients into a group-averaged gradient (group size equalling dataset size) 500 times. Each group-averaged gradient was clustered using the K-means algorithm into *k* = 2,3,…,10 clusters (KM). The results of each clustering were fed into a consensus matrix, one for each clustering solution or *k* (CC). The optimal cluster solution was chosen by calculating the proportion of ambiguous clustering (PAC) in the consensus matrices and Percent Agreement and visualising heatmaps (PAC/HM/PA). The consensus matrix for the optimal cluster solution was then clustered using spectral clustering (SC). Seed-based FC was computed for the resultant clusters.

We used bootstrap aggregation, or bagging, and consensus clustering to select the optimum number of clusters, *k*, to partition each gradient. Bagging improves reproducibility of FC data clustering (Nikolaidis et al., 2020). Here, we implemented bagging by resampling randomly the gradients from the dataset with replacement into a group-averaged gradient multiple times (*n* = 500), with group size equalling dataset size. This was done separately for the exploration and validation samples, and for the dataset as a whole. Each group-averaged gradient was then clustered using the k-means algorithm from scikit-learn (Pedregosa et al., 2011) into *k* = 2,3,…,10 clusters and clustering solutions for each *k* were saved into a matrix and used to build a consensus matrix for consensus clustering (Bellec et al., 2010; Kelly et al., 2012; Ozdemir et al., 2015). Consensus clustering involves building a consensus matrix which quantifies the frequency by which pairs of voxels are placed in the same cluster for each *k* across multiple iterations - in our case across the 500 bagged group-averaged gradients. Here, a consensus matrix for each *k* was built by first building an adjacency matrix **A** of dimension *n* x *n*, where *n* is the number of voxels for the left or right hippocampus, for each group-averaged gradient. The adjacency matrix was such that **A**_ij_ = 1 if voxel *i* and *j* are placed in the same cluster and **A**_ij_ = 0 if they are not placed in the same cluster. The whole set of adjacency matrices for each *k* were then averaged to yield the consensus matrix **C**. All elements in **C** lie in the interval [0,1]. The consensus matrix for a perfectly stable and consistent clustering would contain solely 1s and 0s. The value of *k* can then be chosen from the consensus matrices, under the assumption that the true value of *k* would lead to the most stable clustering (Șenbabaoğlu et al., 2014). The consensus matrix for each *k* can then be clustered to yield the most stable patterns of clustering across group-averaged gradients (Kelly et al., 2012). We clustered the consensus matrices, using the method outlined above, with the spectral clustering algorithm from scikit (Pedregosa et al., 2011). To identify the optimal number of clusters for each gradient, we computed the proportion of ambiguous clustering (PAC) for each consensus matrix. PAC is defined as the fraction of sample-pairs, here voxel-pairs, in the consensus matrix with values falling in the sub-interval (*x*_1_,*x*_2_) η [0,1]. Here, (*x*_1_,*x*_2_) = (0.1,0.9). A low value of PAC indicates stable clustering across group-averaged gradients and therefore a good value of *k* (Șenbabaoğlu et al., 2014). We also computed Percent Agreement across the exploration and validation datasets, by adding the number of voxels that were assigned to the same cluster, after spectral clustering the consensus matrices for each *k*.

### 2.6 Seed-Based Functional Connectivity Analysis

The resulting clusters from spectral clustering were used as seed ROIs in a FC analysis. Whole-brain FC analysis was conducted for each cluster, and each gradient, in each mouse, by quantifying the Pearson Correlation between each cluster and each non-hippocampal ROI in the Allen CCF atlas (**Supp. Figure 1**). For comparison, we also conducted a FC analysis using the DG and CA1-3 as ROIs. Significant group-level correlations (edges) for each cluster were identified by conducting a non-parametric permutation test against zero for the correlations between the cluster and each other ROI across mice and correcting for multiple comparisons using the Benjamini-Hochberg non-negative method with a false discovery rate (FDR) of *p* <0.005.

### 2.7 Hippocampal Gene Co-Expression Gradients

To examine the potential mechanistic underpinnings of the FC gradients, we computed gradients reflecting the main patterns of variation in genomic anatomy across the mouse hippocampus by adapting the gradient analysis method to gene co-expression (GCE) data (see **Figure 3** for a schematic overview of the method). We used the Anatomical Gene Expression Atlas (AGEA; https://mouse.brain-map.org/agea) correlation maps quantifying GCE. The AGEA is an open-access relational atlas of the genetic architecture of the adult C57B1/6J mouse brain. The GCE maps were created by computing voxelwise spatial correlations across GE profiles from *in situ* hybridization (ISH) data from the Allen Brain Atlas (ABA) for 4,376 genes. More detailed methodology for the AGEA can be found in its corresponding paper (Ng et al., 2009). As for the FC gradient analysis, we were interested in the GCE maps for seed voxels in CA1, CA2, CA3 and DG. To acquire the maps, we iterated over the anatomic labels of all seed voxels in a range of x, y and z-coordinates within the AGEA, and downloaded the correlation maps of seed voxels labeled anatomically as ‘CA1’, ‘CA2’, ‘CA3’ or ‘DG’. This yielded 2,786 GCE maps.

**Figure 3:**
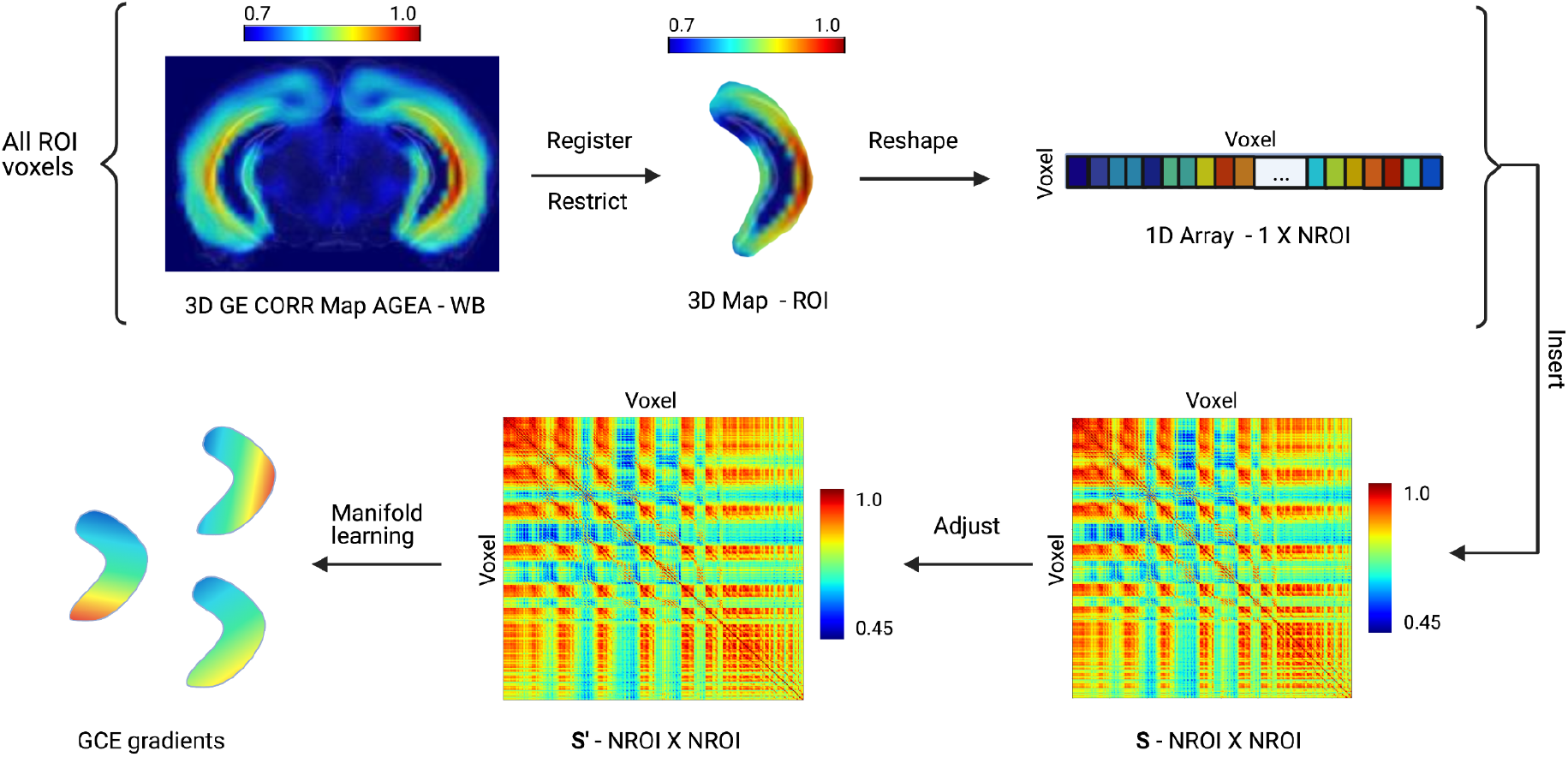
Schematic Overview for the Gene Co-Expression Gradient Analysis Method. Each correlation map from the region of interest (ROI) was registered to the Allen Mouse Brain Common Coordinate Framework (CCFv3) and restricted to contain only voxels within the left or right hippocampus. The rest of the analysis was conducted for the left and right hippocampus separately. Here, an example is given for the right hippocampus. The restricted maps were reshaped into row vectors of length *NROI, NROI* being the number of voxels within the right hippocampus, and inserted into an empty matrix of dimension *NROI* X *NROI*. This resulted in a non-symmetric similarity matrix, **S**, due to the registration of the AGEA correlation maps. **S** was adjusted into a symmetric similarity matrix, **S’**. Manifold learning was then applied to **S’**, yielding a set of gene co-expression (GCE) gradients. Abbreviations in figure, CORR - correlation, WB - whole brain.

The GCE and seed voxel maps were normalized to the Allen Mouse Brain Common Coordinate Framework (CCFv3) (Wang et al., 2020), using linear affine and nonlinear greedy SyN diffeomorphic transformation metric mapping using ANTs v2.1 (picsl.upenn.edu/ANTS). The normalized seed voxel maps were used to build separate left and right hippocampus masks that encompassed all the seed voxels. We then computed the intersection of these seed voxel masks and the hippocampus masks used in the FC gradient analysis. These masks were also used to restrict the GCE maps to include only voxels within the hippocampus, as we were primarily interested in variation in genetic co-expression across the hippocampus rather than the whole brain.

The restricted correlation maps were then used to construct similarity matrices, **S**, one for the left and one for the right hippocampus. The process of normalizing the GCE maps (i.e., correlation matrices) created some minor distortions that did not arise in the processing of the FC data, where normalization occurred before computing correlations. Some correlation maps had the same seed voxel; in that case, the mean of the maps was used for **S**. Other minor distortions in the correlation maps meant the similarity matrices were not symmetric; in that case, the mean of **S** and its transpose **S**^**T**^ was computed to ensure symmetry. These adjusted similarity matrices were highly similar to **S** (see **Figure 3** and **Supp. Figure 14** and **15**).

Manifold learning using Laplacian eigenmaps was then applied to the adjusted similarity matrices in the same manner as the FC gradient analysis to yield GCE gradients. The gradients represent maps characterising the modes of GCE change, where similar values within a gradient mean similar GCE patterns. We chose to retain the three principal gradients for further analysis based on the clearest elbow in Scree Plots from the analysis (see **Supp. Figure 16** and **17**). Elbows in these Scree Plots were not as clear as those obtained for the FC gradient analysis. To compare the GCE and FC gradients, we restricted the FC gradients of the exploration group using the hippocampal mask described above and correlated the gradients using Pearson’s r.

### 2.8 Data, Materials, and Code Availability

The mouse rsfMRI data is available from V. Zerbi upon request. The GCE data are openly available through the AGEA website (https://mouse.brain-map.org/agea). The pre-processing pipeline we used is described extensively in Zerbi et al. (2015). Connectopic mapping is described in detail in Haak et al., 2018, and code for the procedure can be found here: https://fsl.fmrib.ox.ac.uk/fsl/fslwiki/OtherSoftware. FC gradients and maps, GCE maps, code and ROIs used for this study are available through github (https://github.com/BrynjaGunnars/FC_GE_mouse_Hippocampus).

## 3. Results

### 3.1 FC Gradient Robustness across Exploration and Validation datasets

The first three FC gradients were analysed, based on the clearest elbow in a scree plot from the exploration dataset (see **Supp. Figure 2** and **3**). The first two FC gradients were robustly evident at the individual as well as the group level. The third gradient, while still very robust, was not as stable either across hemispheres or individuals. Across the exploration and validation datasets, the first two gradients were almost perfectly correlated (Pearson R = 0.98 and 0.99 for the left and right principal gradients, respectively, and Pearson *R* = 0.99 for the second gradient in both hemispheres), while the third gradient was highly correlated in the right hemisphere (Pearson R = 0.76) and very highly correlated in the left hemisphere (Pearson *R* = 0.92). This suggests that the first two gradients in particular were very robustly identified across samples.

### 3.2 The Principal Gradient of Hippocampal FC Transitions Sharply Along The Long Axis

The principal FC gradient, which represents the dominant mode of FC change, followed the hippocampal long axis in the dorsal-to-ventral direction. In the exploration dataset (*n* = 30), the gradient was relatively uniform dorsally but featured a sharp transition to more ventral regions, suggesting an abrupt change in connectivity. The dorsal region defined by the sharp transition is hereafter referred to as the dorsal gradient region (DGR) and the region ventral to it as the ventral gradient region (VGR). To our knowledge, the DGR does not correspond to any anatomical subdivision of the hippocampus. Ventrally to the sharp transition, the gradient exhibited a smaller and more gradual change (see **Figure 4**). In the validation dataset (*n* = 20), the gradient of the left hippocampus featured a more gradual transition from dorsal to ventral hippocampus (see **Supp. Figure 4**).

**Figure 4:**
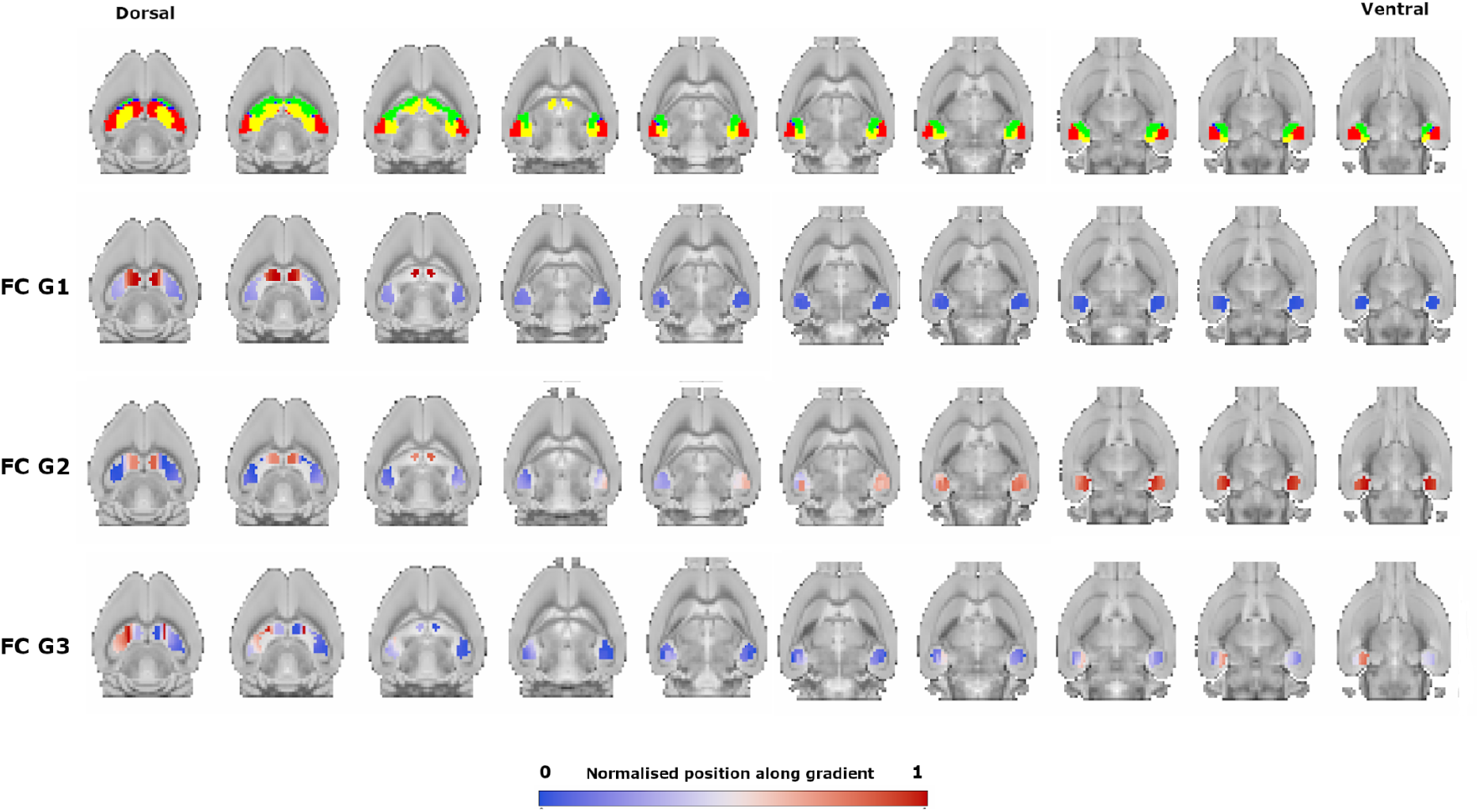
FC Gradients. The figure shows the topography of the first three FC gradients (FC G1-G3) in the exploration dataset. The gradient analysis was conducted on the left and right hippocampus separately. They are presented together here to enable comparison across hemispheres. The images are horizontal slices of a mouse brain. The first row shows the location of the hippocampus subfields in the mouse brain in the dorsal-to-ventral direction, where CA1 is red, CA2 is blue, CA3 is green and the DG is yellow. The second row shows the topography of the first FC gradient. Note that gradient values are unitless. Similar colours indicate similar connectivity fingerprints. The gradient follows the hippocampal long axis and features a sharp transition from dorsal to ventral hippocampus. In the second FC gradient in the third row, the DGR is distinct and the gradient transitions gradually along the hippocampus long axis in the VGR. The third FC gradient is in the fourth row and features a sharp line dorsally at one extreme of the gradient and shifts towards the other extreme towards the lateral most portion of the hippocampus. When comparing separate gradients, the topography is the main feature, not different colours.

To further examine the nature of connectivity change along the hippocampal long axis, we plotted gradient value as a function of the euclidean distance from the most dorso-medial location of the hippocampus (see **Figure 5**). The graph reflected the same pattern of abrupt change described above. Gradient values fell sharply from the DGR to the VGR, but within the VGR, gradient values changed more gradually.

**Figure 5:**
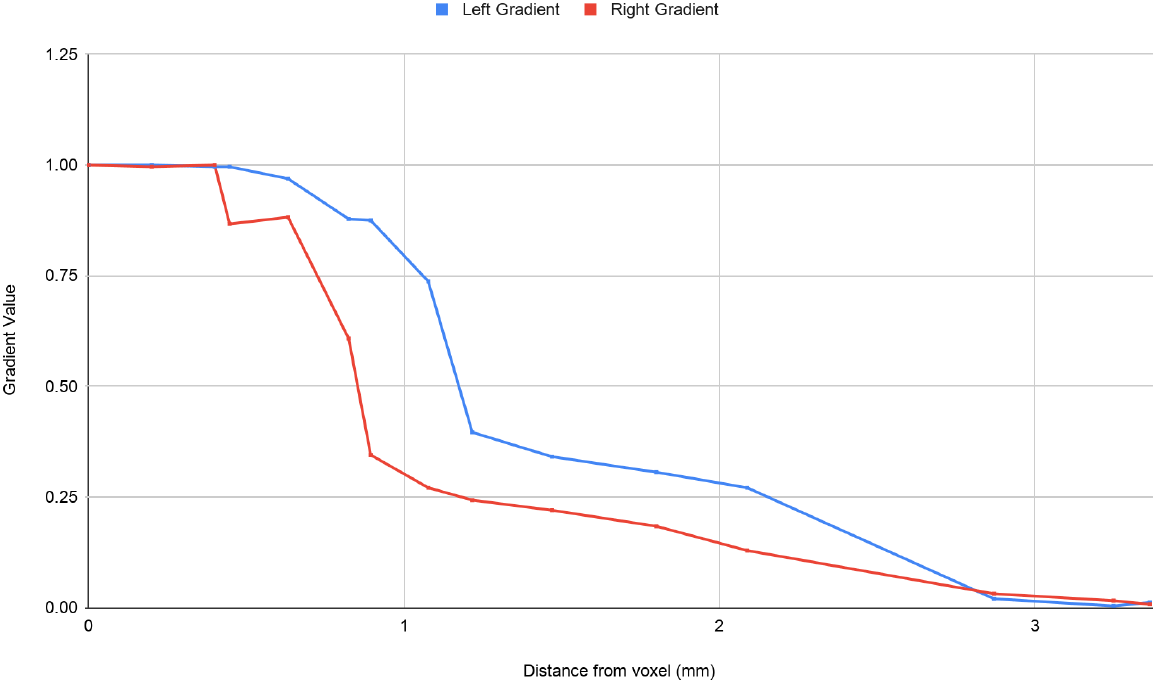
Principal FC Gradient Long Axis Connectivity Change. Distance, in mm, from the most dorso-medial location of the hippocampus plotted against gradient value in the exploration dataset. The principal FC gradients in the left and right hippocampus change sharply from the DGR to the VGR. The gradients vary less in ventral regions and more gradually.

Given the apparently sharp transition indicated by the gradient, we applied consensus clustering and found that the principal gradient of the left hippocampus was best characterised by two clusters (*PAC* = 0.031, Percent Agreement > 95%, see **Supp. Figure 6** and **7** for heatmaps and *PAC* of all cluster solutions) while the principal gradient of the right hippocampus by four clusters (*PAC* = 0.058, Percent Agreement > 95%, see **Supp. Figure 6** and **7** for heatmaps and *PAC* of all cluster solutions). The DGR was reliably separated into its own cluster in both hemispheres. **Figures 6, 7** and **Supp. Figure 5** show the FC maps for each of the left and right principal gradient clusters and for comparison, for the hippocampus subfields. See **S2** in the **Supplementary Material** for comparison between the FC maps of the left and right gradient clusters.

**Figure 6:**
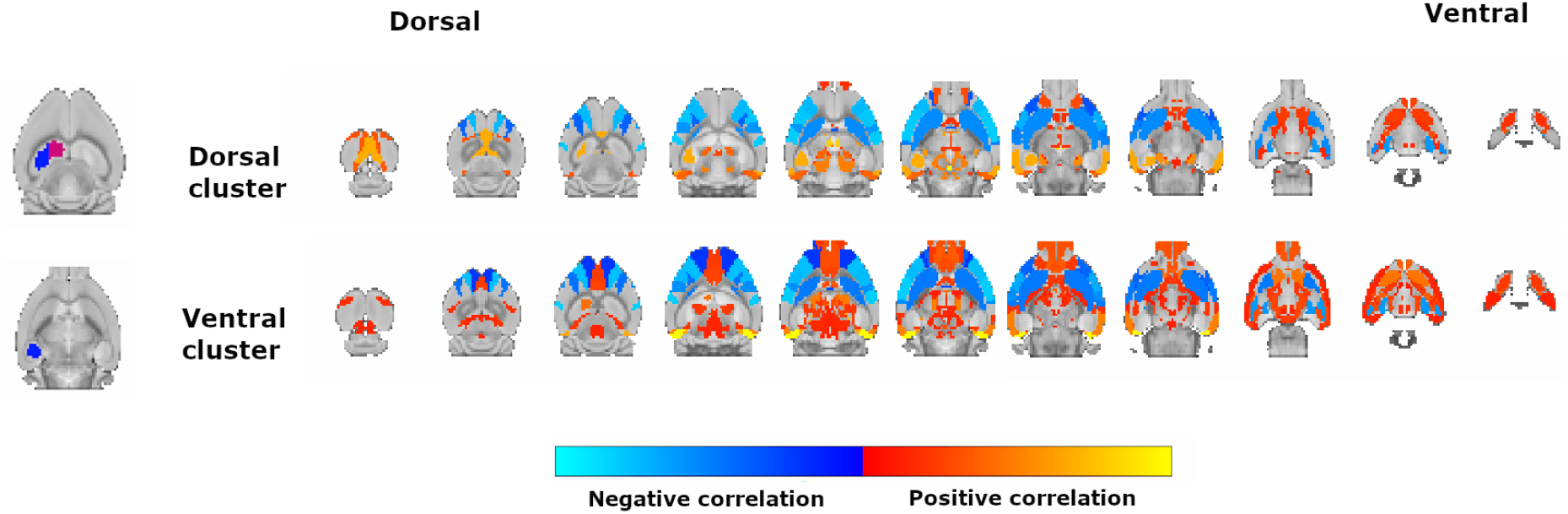
Significant Correlations of the Left Principal FC Gradient. The figure shows the different patterns of correlations across the gradient clusters of the left principal FC gradient. On the left are the clusters resulting from consensus clustering in the dorsal-to-ventral direction from top to bottom. Each row shows significant correlations (Pearson R) for a gradient cluster. Positive correlations are on the red spectrum, where a more yellow colour signifies a stronger positive correlation for each cluster. Negative correlations are on the blue spectrum, where a lighter blue signifies a stronger negative correlation.

**Figure 7:**
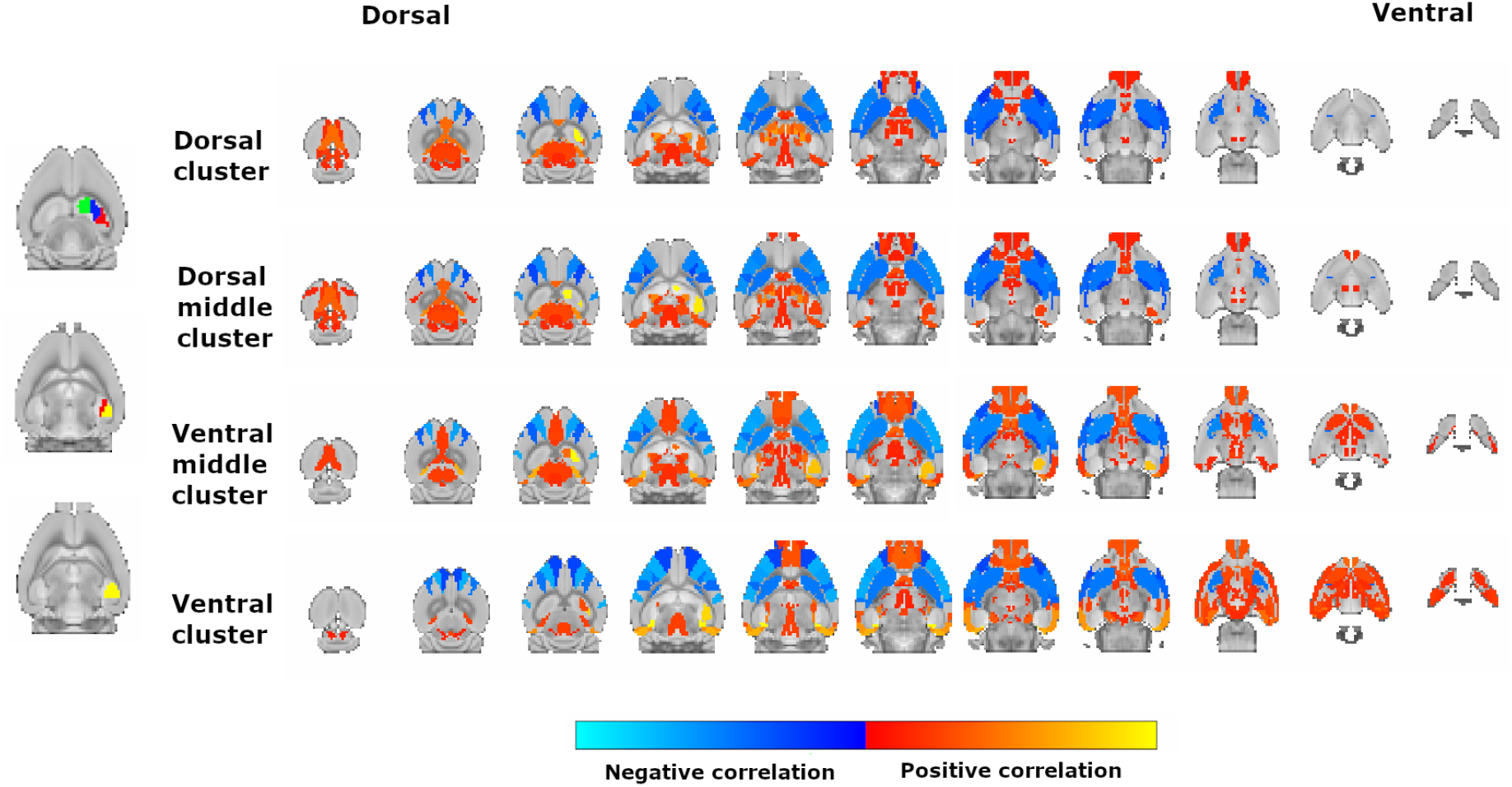
Significant Correlations of the Right Principal FC Gradient. The figure shows the different patterns of correlations across the four clusters of the right principal FC gradient.

Dorsal regions of the first principal gradient were significantly positively correlated with the retrosplenial cortex and showed sparse FC with prefrontal areas, while ventral regions were more strongly correlated with prefrontal regions, such as the prelimbic, infralimbic and anterior cingulate cortex and only dorsal regions within the VGR in the right hippocampus were correlated with the posterior parietal association cortex. FC with olfactory regions increased ventrally. Subcortically, FC with superior colliculus decreased ventrally, while FC with amygdalar and hypothalamic nuclei increased. In the right gradient, FC with anterior thalamic nuclei decreased ventrally as well.

Common patterns of FC were also evident along the gradient (i.e., across clusters), including FC with thalamus, entorhinal and orbitofrontal (excl. agranular insular) cortex, septal nuclei, presubiculum, parasubiculum and postsubiculum. Interestingly, negative FC was highly similar across clusters, and all three gradients, where somatosensory, motor and agranular insular areas were consistently segregated from the hippocampal networks.

Comparing **Figure 6** and **Figure 7** with **Supp. Figure 5**, it is clear that the gradient clusters reveal transitions in FC patterns across the hippocampus that are not evident when examining the FC of the hippocampus anatomical subfields. Patterns of FC for the hippocampus subfields were on the whole very similar, including amygdalar, olfactory, thalamic, hypothalamic, prefrontal cortical and posterior cortical regions and regions of the rodent equivalent of the Default-Mode Network (DMN), i.e., the posterior parietal, anterior cingulate, orbitofrontal and retrosplenial cortex (Stafford et al., 2014). The main differences between hippocampus subfields were stronger FC with midbrain and hindbrain areas of the CA fields in comparison to the DG.

### 3.3 The Second Gradient Follows the Hippocampal Long Axis in the VGR

The second gradient also captured connectivity variation along the hippocampal long axis, though it differed from the principal gradient. The DGR was distinct in this gradient as well, where the gradient changed sharply to the VGR. In the VGR however, the gradient followed the long axis in a gradual manner (see **Figure 4**).

As might be expected for a gradient without clear boundaries or transition points, consensus clustering of the second gradient did not reveal an optimal solution, as indicated by the high *PAC* for all cluster solutions compared to the principal gradient and heatmaps. For our purposes of examining the broad features of FC that change across the gradient, we examined the FC associated with the six-cluster solution (*PAC* = 0.41 and 0.40 for the left and right gradient, respectively, and Percent Agreement > 95%, see **Supp. Figure 8** for heatmaps; **Supp. Figure 9** for *PAC* of all cluster solutions; **Figure 8** and **Supp. Figure 10** for FC maps).

**Figure 8:**
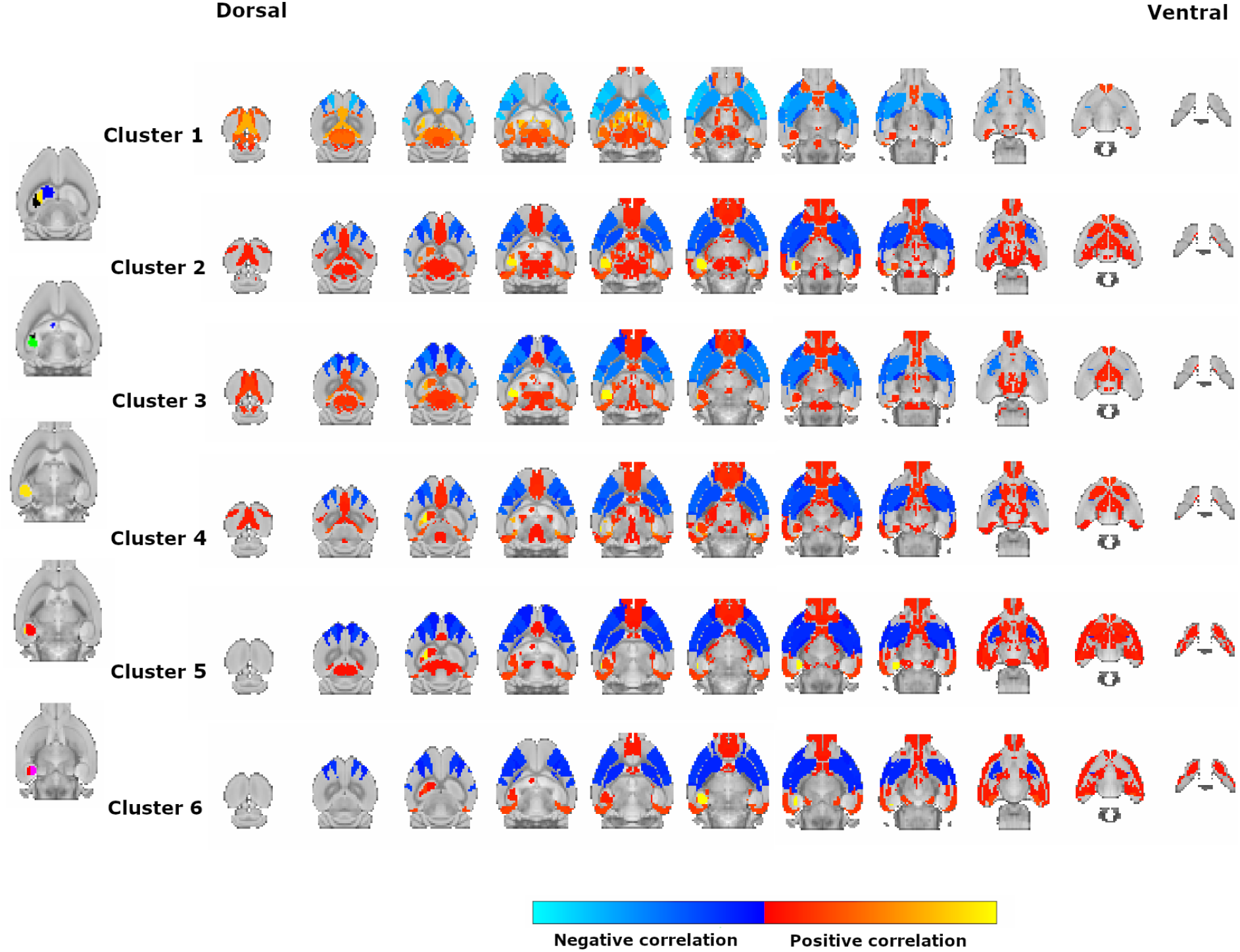
Significant Correlations of the Left Second FC Gradient. The figure shows the different patterns of correlations across the gradient clusters for the second FC gradient of the left hippocampus (cluster 1 - blue, cluster 2 - yellow, cluster 3 - black, cluster 4 - green, cluster 5 - red, cluster 6 - pink)

The DGR was reliably clustered in both hemispheres, and therefore had highly similar patterns of FC as the DGR clusters of the first gradient. Although the variation in FC along the VGR of the left second gradient was more subtle compared to the differences between the DGR and VGR, we found some interesting patterns. The main pattern we found was an increase in significant FC with amygdalar nuclei when moving from dorsal regions of the VGR to ventral. Additionally, the ventral most portion of the VGR was functionally connected with the infralimbic and prelimbic cortex, but otherwise had relatively sparse prefrontal, thalamic and hypothalamic correlations compared to the other clusters. Cortical connectivity was more similar for other regions. However, FC with retrosplenial, anterior cingulate and posterior parietal association cortex were more prevalent in the dorsal half of the VGR. Examining similarities across clusters, we found the same general pattern of FC across clusters that was observed for the first gradient: a core common network comprising thalamus, entorhinal and orbitofrontal (excl. agranular insular) cortex, septal nuclei, presubiculum, parasubiculum and postsubiculum.

### 3.4 The Third Gradient Follows the Medial-to-Lateral axis of the Ventral Hippocampus

The third gradient diverged from the pattern suggested by the first two, in that it varied along the medial-to-lateral axis (see **Figure 4**), although the DGR was distinct in this gradient as well. This gradient was not as stable as the first two, and, while still very robust, showed greater variation across hemispheres, individual mice, and samples.

The gradients were divided into two clusters through consensus clustering (*PAC* = 0.20 and 0.24 for the left and right gradients, respectively, Percent Agreement > 95%, see **Supp. Figure 11** and **12** for heatmaps and *PAC* of all cluster solutions), though in each hemisphere both clusters were split in two. See **Figure 9** and **Supp. Figure 13** for the significant correlations of the left and right gradients. The significant positive FC was largely shared across clusters, with prevalent FC with amygdala, olfactory areas, prefrontal cortex, thalamus, striatum, hypothalamus and midbrain regions. The two clusters differed mainly in terms of their thalamic and midbrain correlations (see **Figure 9**). The dorsal/middle cluster exhibited FC with the central lateral nucleus, mediodorsal, posterior limiting and ventral posteromedial nucleus of the thalamus while the dorsal/ventral cluster exhibited strong FC with more anterior thalamic nuclei and the ventral anterior-lateral and posterior complex of the thalamus. The dorsal/middle cluster had stronger FC with midbrain areas, including midbrain reticular, red and interpeduncular nucleus and the ventral tegmental area while the dorsal/ventral cluster exhibited stronger FC with the reticular part of the substantia nigra and the sensory related regions of the superior colliculus.

**Figure 9:**
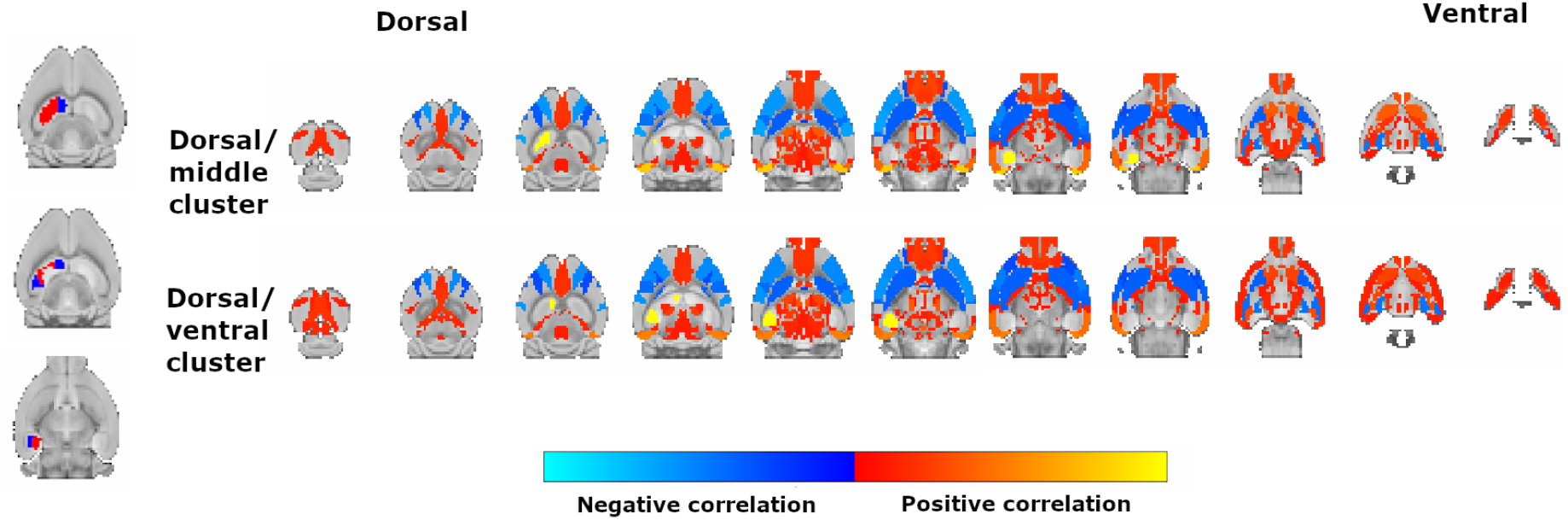
Significant Correlations of the Left Third FC Gradient. The figure shows the different patterns of correlations across the gradient clusters of the left third FC gradient.

### 3.5 Contribution of Gene Co-Expression Gradients to FC Gradients

#### 3.5.1 Intrinsic Dimensionality of the AGEA Correlation Maps

We were interested in how genomic anatomy contributes to the FC gradients we uncovered, especially given the novel patterns of functional organization we obtained for the gradients relative to the anatomical subdivisions. The elbow in the GCE scree plots was not as clear as in the FC scree plots, and the relative eigenvalue of the principal gradients was smaller compared to the principal gradients of FC (0.34 and 0.39 for the left and right GCE gradients compared to 0.63 and 0.66 for the left and right FC gradients) (see **Supp. Figure 16** and **17** for the scree plots of the GCE gradients). This indicated that the first GCE gradient captured less of the variation in the GCE data compared to the first FC gradient, and that the GCE variation might be distributed across more gradients.

#### 3.5.2 Gene Co-Expression Gradients Highlight Variation in the Ventral Hippocampus

The three principal gradients mainly describe GCE variation within the intermediate and ventral thirds of the hippocampus. The gradients were similar across hemispheres. The principal gradient of gene co-expression described variation between the hippocampus subfields, mainly in more intermediate and ventral areas of the hippocampus. Dorsally, a variation was seen between the DG and the CA fields. The second gradient described variation ventrally, mainly between an anterior and posterior region within the CA3 field laterally and between the CA3 and the DG. The third gradient described GCE variation in a similar region as the second gradient. However, the main variation was in the lateral to medial direction (see **Figure 10** for the GCE gradients).

**Figure 10:**
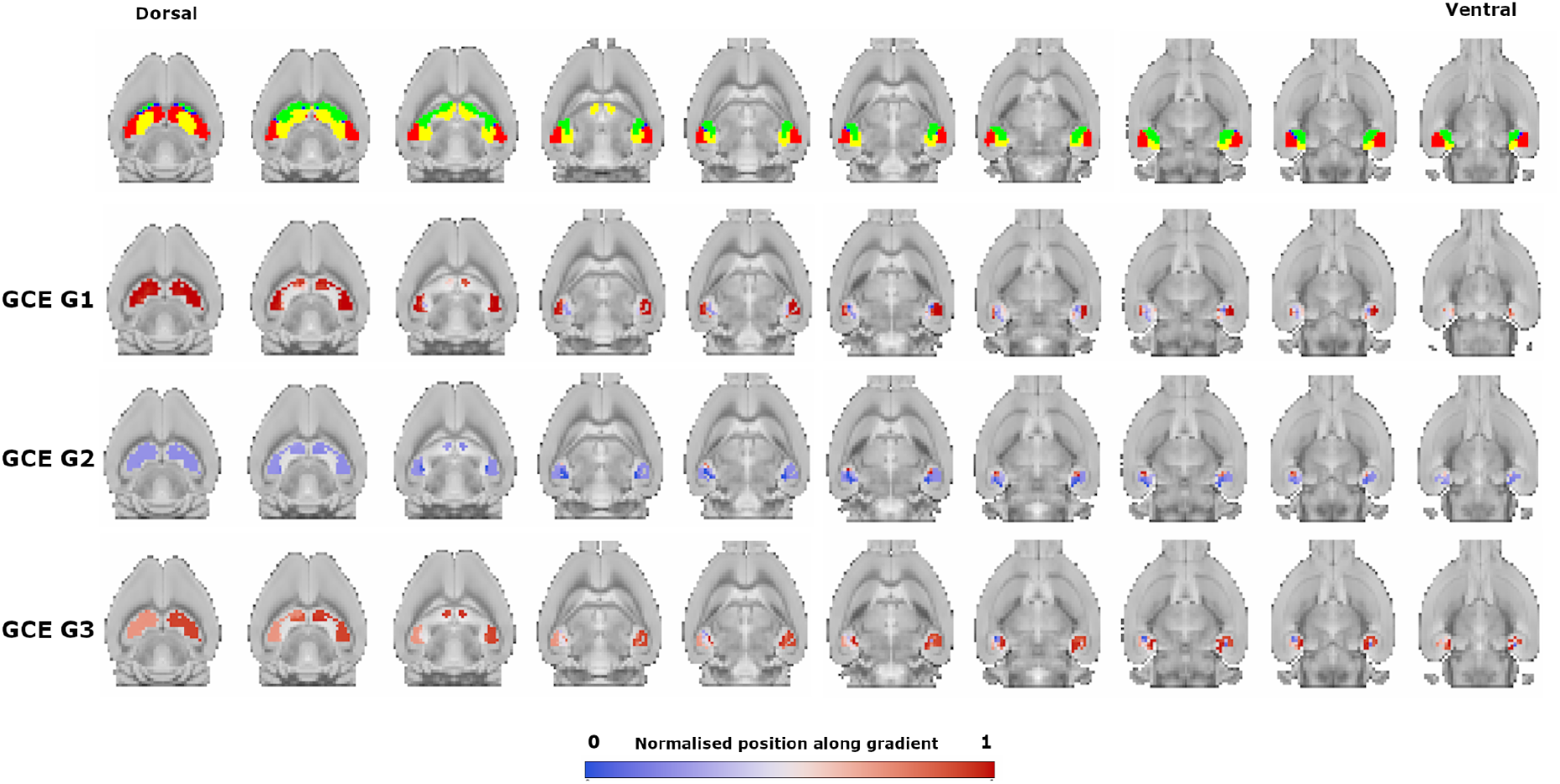
GCE Gradients. The first row shows the location of the hippocampus subfields in the mouse brain in the dorsal-to-ventral direction, where the CA1 is red, the CA2 is blue, the CA3 is green and the DG is yellow. The second row shows the topography of the first GCE gradient (GCE G1). Similar colours indicate similar gene co-expression profiles. The gradient describes GCE variation between the subfields, mainly in more intermediate and ventral areas of the hippocampus. Dorsally, the gradient varies between the DG and CA fields. The second GCE gradient (GCE G2) in the third row describes GCE variation ventrally, mainly between an anterior and posterior region within the CA3 field laterally and between the CA3 and the DG. The third GCE gradient (GCE G3) in the fourth row describes GCE variation in a restricted region ventrally in the lateral-to-medial direction.

Comparing the GCE gradients with the FC gradients visually, we did not see a strong similarity across modalities (see **Figure 11**). The FC gradients mainly capture large-scale variation along the hippocampus that does not appear to be constrained by the hippocampus anatomical subfields. A sharp distinction between the DGR and VGR is also evident. In contrast, the GCE gradients describe a variation in more restricted regions where differences between and within subfields seem to play a bigger role in the gradient topography. Moreover, the GCE gradients mainly capture variation ventrally and the DGR is not evident in any of the gradients.

**Figure 11:**
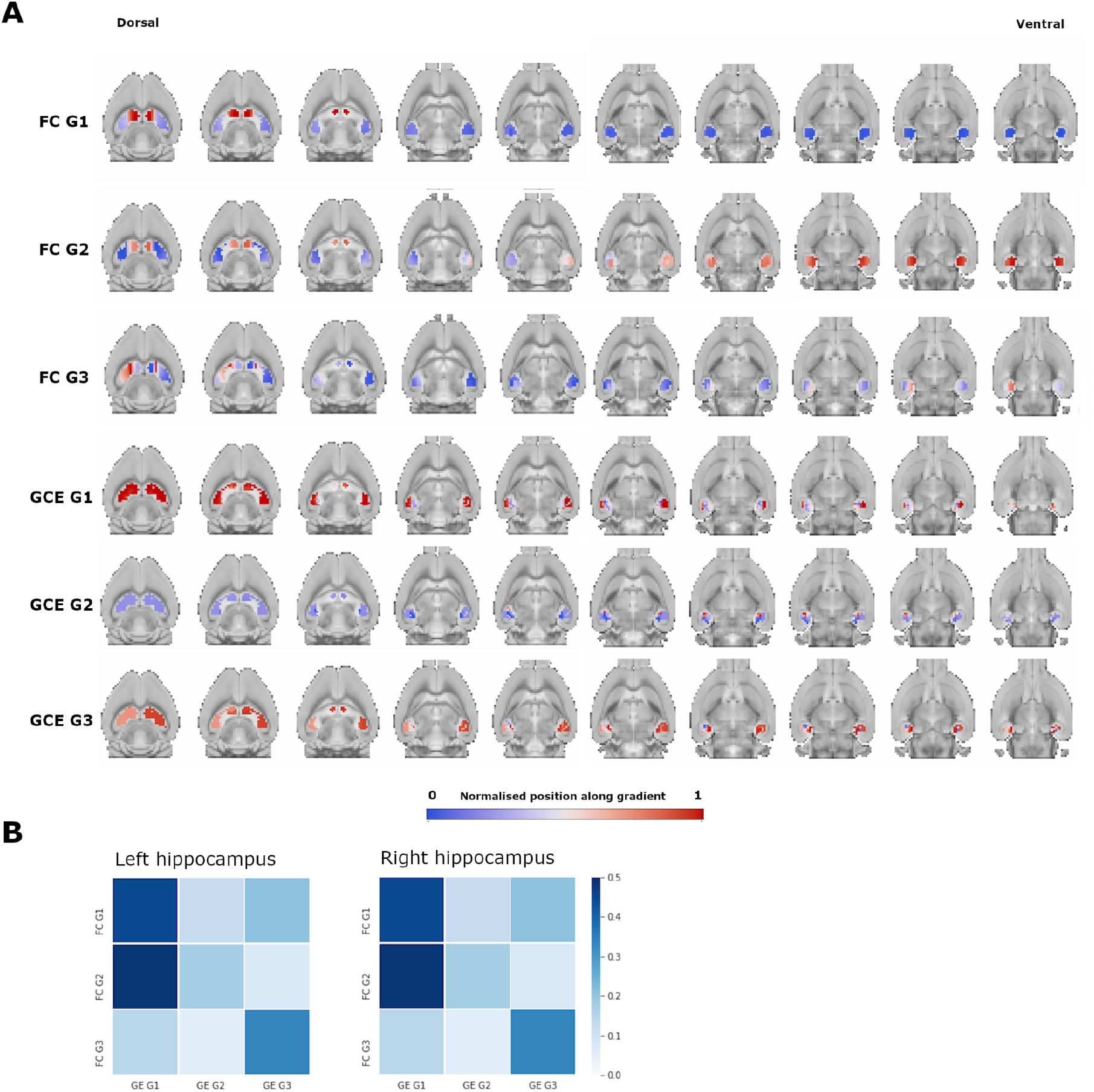
Comparison Between the FC Gradients and the GCE Gradients. Figure allowing comparison between the FC and GCE gradients. A shows the topography of the first three FC gradients (FC G1-G3) and the GCE gradients (GCE G1-G3). B shows the correlation matrices for the correlations between the FC and GCE gradients of the left and right hippocampus. A darker blue colour indicates a stronger correlation, where the maximum is set at *R* = 0.5 for ease of comparison.

We quantified the cross-modality similarity by computing the spatial correlation (Pearson *R*) between the gradients from the two modalities. Weak-to-moderate relationships were observed (see **Figure 11B**), with the strongest relationship being observed between the first GCE gradient and the first FC gradient (*R* = 0.29, *p* = 8.9 × 10^−9^, and *R* = 0.45, *p* = 1.2 × 10^−21^, in the right and left gradients), between the first GCE gradient and the second FC gradient (*R* = 0.43, *p* = 1.3 × 10^−18^, and *R* = 0.49, *p* = 2 × 10^−25^, in the right and left gradients), and between the third GCE gradient and the third functional connectivity gradient (*R* = 0.34, *p* = 4.7 × 10^−12^). The second GCE gradient did not show strong correspondence with any FC gradient. Absolute correlation values are reported as the sign of a correlation does not matter in this context. All *p*-values were significant, even after correction for multiple comparisons.

## 4. Discussion

### 4.1 Summary of Findings

Despite being one of the most commonly studied brain structures, the functional organisation of the hippocampus remains incompletely understood. We examined the functional organisation of the hippocampus in mice by uncovering the main axes of variation in hippocampal whole-brain FC, and examining correspondence with patterns of variation in gene co-expression across the hippocampus. We chose a novel analysis method, gradient analysis, for its ability to capture sharp, as well as gradual, variation in data and to tease out multiple, potentially overlapping principles of variation.

The FC gradients were robustly evident at the individual and group level. The two main FC gradients revealed variation principally along the hippocampal long axis, as we expected. Unexpectedly however, the first gradient effectively divided the hippocampus into two distinct regions, the dorso-medial extreme (DGR) and the rest of the hippocampus (VGR). The second gradient revealed a gradual change in connectivity along the long axis in the VGR. The third gradient differed from the first two and showed variation in the medial-to-lateral direction in the VGR. Consensus clustering and analysis of FC patterns driving the gradients revealed both established and less well-known patterns of connectivity.

To examine whether these FC gradients were related to the underlying genomic anatomy, we adapted gradient analysis to gene co-expression (GCE) data, and uncovered the main patterns of GCE variation within the hippocampus. The GCE gradients did not reveal a long axis gradient but a more anatomically constrained and restricted variation in GCE compared to FC, whereby the first gradient captured the anatomical subfields (DG and CA1-3) while the second and third gradients highlighted GCE variation within a small ventral region. These data suggest that the main organisational patterns of FC within the hippocampus are unlikely to be primarily driven by the main variations in GCE, or that there is a more complicated relationship between genomic anatomy and FC within the hippocampus.

### 4.2 Hippocampal Functional Organisation Revealed By Gradient Analysis

Our primary finding was that hippocampal long axis functional organisation is characterised by both sharp functional segregation dorsally and gradual changes in functional organization ventrally. The DGR appeared as a distinct zone in all three primary gradients and was reliably separated by cluster analysis of the first two gradients, which adds further support to this notion. These findings are new and help shed light on disagreements in the literature about hippocampal functional organisation - specifically, they suggest that hippocampal organisation exhibits both sharp discontinuities and gradual change. This is in line with more complex models of hippocampal organisation, such as the model proposed by Strange et al. (2014), whereby long axis functional gradients are superimposed onto discrete functional domains.

The sharp connectivity change between the dorsal and ventral hippocampus revealed by the principal gradient was surprising given recent gradient analyses of human hippocampus, which reported a gradual transition in hippocampal-cortical FC along the long axis (Przeździk et al., 2019; Vos de Wael et al., 2018). The DGR contrasted with the rest of the hippocampus mainly in terms of its strong functional connectivity with the retrosplenial cortex but relatively sparse functional connectivity with other cortical regions. The DGR was also functionally connected with anterior thalamic nuclei and the superior colliculus, regions which are implicated in exploration, movement and orientation (Vogel et al., 2020). These patterns suggest that DGR may play a distinct role in aspects of spatial cognition, a hypothesis that should be examined in further work. Examining homology of FC across species is somewhat complicated by the challenge of matching corresponding cortical regions. We did find several similarities across species, such as greater connectivity between the ventral hippocampus and temporolimbic regions (e.g., perirhinal cortex), and between the dorsal hippocampus (excluding the DGR) and anterior cingulate cortex (implicated in attention) and posterior parietal cortex (involved in sensory processing) (Lyamzin and Benucci, 2019; Han et al., 2002). Unlike gradient analyses of the human hippocampus, however, we did not find stronger connectivity of the dorsal hippocampus to visual cortex or attention regions, or of the ventral hippocampus to somatomotor or default network regions. Although the hippocampus is a phylogenetically preserved structure across mammalian species, there are important differences in structure and connectivity (Andersen et al., 2007; Bergmann et al., 2016). The different pattern of results obtained here may reflect species differences, the effects of anaesthesia, or other methodological differences; future studies should attempt to resolve these influences.

The functional differences along the VGR in the second gradient were more subtle than the sharp distinction between the DGR and VGR. Cortical functional connectivity was strongest in dorsal and intermediate regions, with functional connectivity to e.g., anterior cingulate cortex (implicated in e.g. memory processing, attention and cognitive control), and preferential functional connectivity to retrosplenial and posterior parietal cortex dorsally (Brockett et al., 2020; Frankland et al., 2004; Han et al., 2003). All regions of VGR shared functional connectivity with the infralimbic, prelimbic and orbitofrontal cortex. Additionally, amygdalar functional connectivity increased from dorsal to ventral regions. Interestingly, the ventral-most region had little connectivity with hypothalamic nuclei. These differential patterns of FC are potentially novel and their significance merits further examination. While the gradient may reflect a dorsal-ventral functional transition from spatial navigation, cognition, and learning toward affect and fear, it might also point to more subtle functional distinctions, such as differential involvement in certain types of memory, with more involvement in e.g., spatial memory dorsally and emotional memory ventrally (Ritov et al., 2014).

Our third FC gradient revealed a medial-to-lateral transition in the VGR. Neither the follow-up FC analysis of the gradient or the GCE analysis revealed a clear basis for this gradient. In humans, Vos de Wael et al. (2018) found a second gradient that also followed a medial-to-lateral direction. This gradient was significantly correlated with intracortical microstructure, indicating interactions between tissue microstructure and hippocampal function. It is therefore possible that our third gradient is related to the underlying microstructure; additional multimodal data could shed light on this. Taken together, the gradients we described suggest the multiple roles of the hippocampus are subserved by both discrete and overlapping principles of functional organisation. Such complex organisation might enable efficient wiring in the hippocampus, e.g., to efficiently integrate multimodal information reaching it for episodic memory formation (Andersen et al., 2007; Zemla and Basu, 2017; Haak and Beckmann, 2020).

The benefits of the gradient approach were revealed by comparing the functional connectivity of the hippocampus anatomical subfields to the patterns obtained across clusters obtained for the first two FC gradients. The FC analysis of the subfields showed distributed and largely undifferentiated functional connectivity, while FC analysis of the gradient clusters revealed substantial differences in functional connectivity across the hippocampus. This is in line with other studies that have underlined the importance of using functionally rather than anatomically defined regions of interest for analysis of fMRI data, since the former capture transitions and distinctions within anatomically homogenous ROIs (e.g., Harrison et al., 2014; Ren et al., 2019). Our findings support this perspective, while also emphasising the existence of both sharp discontinuities and gradual change in functional organisation.

### 4.3 Gene Co-Expression Gradients

What role might gradients in hippocampal GCE play in the functional organisation of the hippocampus? We found possible subfield differences, where GCE variation was less prominent within CA1 compared to CA3, which is in line with earlier suggestions that CA1 and CA3 differ in GCE variation (Cembrowski et al., 2016; Dong et al., 2009). The main long axis pattern in GCE was the more prominent GCE variation in intermediate and ventral regions within the hippocampus. Given previous research on hippocampal GCE variation finding distinct molecular domains along the long axis, we expected to find a more prominent long axis pattern (Dong et al., 2009; Fanselow and Dong, 2010; Thompson et al., 2008).

The GCE gradient analysis indicated the main variation in hippocampal genetic organisation is guided by different principles than the main patterns of hippocampal FC variation. Aligning with that observation, Huntenburg et al. (2021) found that the second and third principal components of cortical GE in mice were significantly correlated to the fifth and third gradients of cortical FC, respectively. The third FC gradient traversed between audiovisual and somatomotor regions and the fifth traversed between sensory and transmodal regions.

Studies examining differentially expressed genes within the hippocampus have found highly represented functional categories which could be important for the establishment and maintenance of FC, such as genes involved in cell adhesion and axon guidance (Lein et al., 2007; Thompson et al., 2008). Additionally, the larger role of the ventral hippocampus in response to stress is reflected at the level of GE where the ventral hippocampus is particularly sensitive to stress (Fanselow and Dong, 2010; Floriou-Servou et al., 2018). Thus, differential gene expression within the hippocampus may influence the FC along the long axis. It is possible a GCE variation along the long axis of the hippocampus exists, gradual or parcellated, but is driven by a small number of genes or a specific subset of genes that is not captured in our analysis. A recent study examining the human cerebellum combined the Allen Human Brain Atlas (AHBA) transcriptome data with a cerebellar functional parcellation atlas and found that a majority of the 443 network specific genes were specific in the limbic (n = 170) and visual (n = 221) networks (Wang et al., 2021). Another recent study found that the location of human tissue samples extracted along the hippocampus long axis could be predicted within 2 mm using the expression pattern of less than 100 genes (Vogel et al., 2020), while our analysis included 4376 genes mapped in the AGEA dataset.

### 4.4 Limitations

Some limitations should be noted. We conducted gradient analysis on a dataset with one fMRI run from a single laboratory. Our findings should be validated in additional multisite data. Additionally, the connectopic mapping method is generally utilised for human data. Although the data acquisition and preprocessing were optimised, challenges specific to mouse fMRI such as physiological noise, signal-to-noise ratio, and anaesthesia could impact our results and their comparability to human hippocampus gradients (Hoyer et al., 2014). Anaesthesia has been shown to affect the neurovascular coupling fMRI relies on to assess neural activity (Reimann and Niendorf, 2020). However, a multi-center comparison of the organisation of mouse FC networks across levels of anaesthesia found that, while there is some effect of anaesthesia, networks were highly robust to anaesthetic protocols (Grandjean et al., 2020). The robustness of the gradients and FC networks identified, along with their correspondence with observations in awake humans, supports our interpretation that these are reliable networks that were robust to the anaesthetic protocol. Combined medetomidine-isoflurane anaesthesia may provide the most specific and reliable FC networks, and we recommend that as an option for further research (Grandjean et al., 2020; Reimann and Niendorf, 2020). Awake imaging is also a possibility, but the influence of stress and arousal would have to be considered, particularly for the examination of hippocampal networks.

### 4.5 Conclusion

Examining the functional organisation of the hippocampus in mice using data-driven methods revealed a complex landscape whereby some functions might be clearly segregated along the hippocampal long axis, while others exhibit a more gradual distribution. While we sought to maximise comparability between the FC and GCE modalities, we did not find compelling evidence of organisational convergence. Nonetheless, our study highlights the value of conducting gradient analyses in mice in future studies. Gradient analysis can capture a complex functional organisation, and the insights that it offers can be combined with data modalities only obtainable in rodents (e.g., involving invasive or terminal experiments) to provide mechanistic insights that are not possible in humans. Our findings thus constitute another step in the translational bridge from rodent to human studies and towards constructing a more complete picture of the principles and determinants of brain functional organisation.

## ACKNOWLEDGEMENTS

The author would like to thank Roselyne Chauvin and Koen Haak for kindly answering inquiries about the Congrads method.

**Figure 1, 2**, and **3** were created with BioRender.com Anaconda Software Distribution, 2020.

## Supplementary Material

### S1 Scree Plots

The gradients uncovered through gradient analysis are eigenvectors, each belonging to one eigenvalue, and are ordered according to importance. We chose to investigate the three principal connectopic maps further, based on results from a scree plot (See **Supp. Figures 2** and **3**). Scree plots have been applied to evaluate intrinsic dimensionality of fMRI data and are a part of the gradient analysis toolbox BrainSpace (Vos de Wael et al., 2020,2018). Scree plots are used to visualise results from dimensionality reduction techniques, mainly principal component analysis (PCA). Scree plots can also be used for Laplacian eigenmaps. The relative eigenvalues are plotted against eigenvalue number and a decision of how many eigenvectors or gradients to retain is based on a specific cut-off value or the clearest elbow in the plot (C.R. Rao et al., 2011). In our case the relative reciprocal eigenvalues (reciprocal eigenvalue divided by the total reciprocal eigenvalue sum) from a group gradient analysis on 10 mice were plotted against eigenvalue number for left and right hippocampus separately. The reciprocal eigenvalue is used instead of the original eigenvalue as the lowest eigenvalue explains the largest variance in the data, as opposed to methods such as PCA where the highest eigenvalue explains the largest variance.

### S2 FC Patterns of Left versus Right Gradient Clusters

The spatial distribution of FC was on the whole similar across the left and right gradient, differing mainly in terms of thalamic and midbrain correlations (see **Figure 6** and **7**). The principal gradient of the left hippocampus was significantly positively correlated to the red nucleus while the right was correlated to more anterior thalamic nuclei and the inferior colliculus. The DGR of the left gradient was correlated to the posterior limiting nucleus of the thalamus, the medial geniculate complex and the midbrain reticular nucleus while the DGR of the right gradient was correlated to the mediodorsal nucleus of the thalamus. The VGR of the left gradient was correlated to the submedial and the ventral posteromedial nucleus of the thalamus while the VGR of the right gradient was correlated to the lateral habenula, the intralaminar nuclei of the dorsal thalamus, the superior colliculus and the substantia nigra. Other differences were e.g. the DGR of the left gradient was correlated to the frontal pole of the cerebral cortex, the somatosensory area for the trunk and the posterior amygdalar nucleus and the DGR of the right gradient to the main olfactory bulb. Additionally, the VGR of the right gradient was correlated to the retrosplenial cortex.

### S3 Supplementary Figures

**Supplementary Figure 1:**
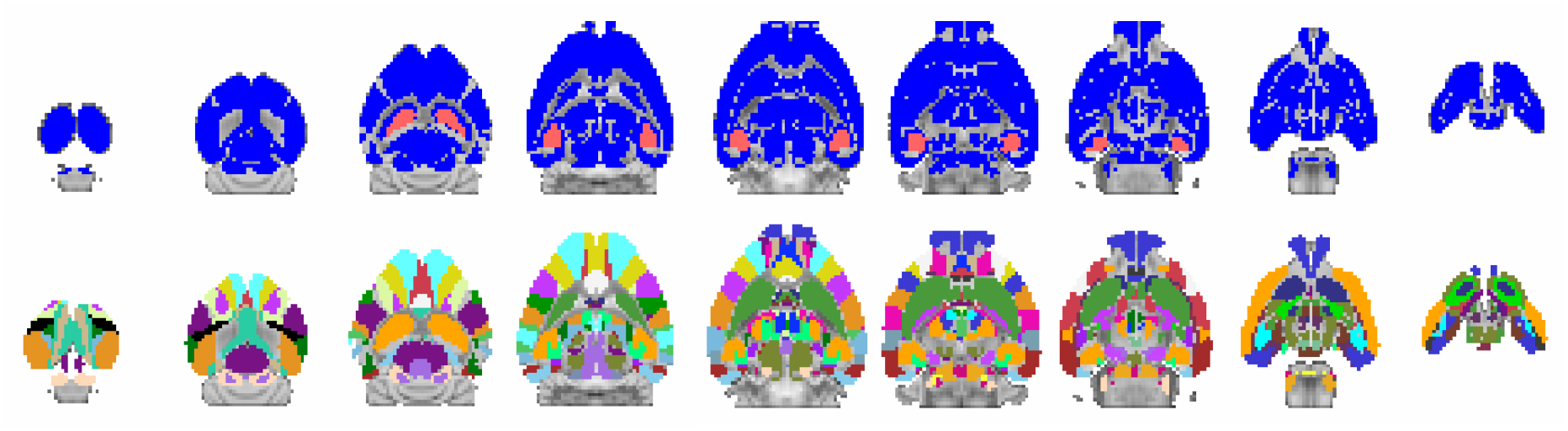
FC Analysis Masks and ROIs. The top row shows the hippocampus masks, in pink, and the whole-brain mask, in blue, used in the gradient analyses overlaid on the template space. The bottom row shows the ROIs in the Allen Common Coordinate Framework (V3, http://help.brain-map.org/download/attachments/2818169/MouseCCF.pdf).

**Supplementary Figure 2:**
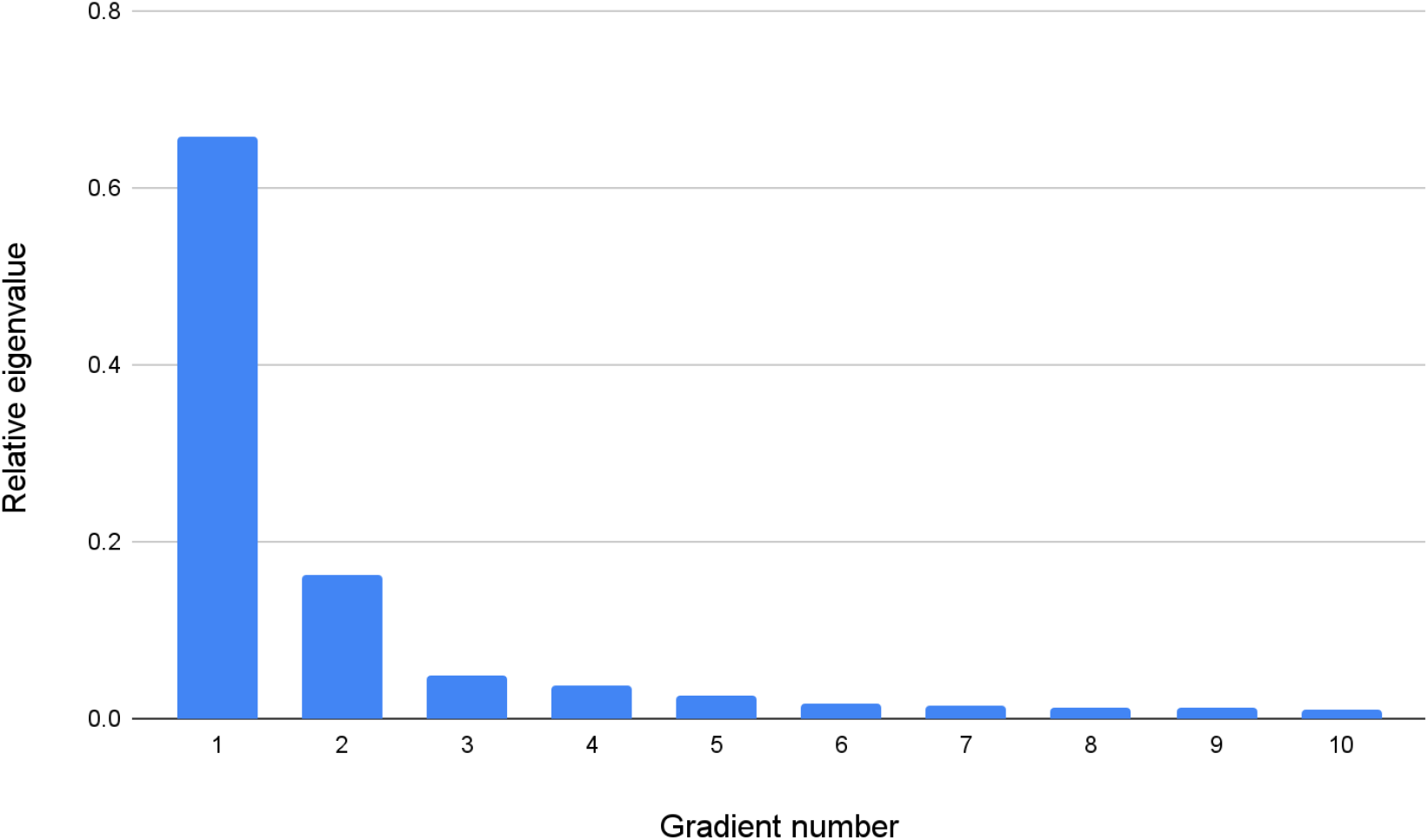
Scree Plot for Left Hippocampus FC Gradients in Exploration Dataset. Relative eigenvalue (reciprocal of eigenvalue divided by total eigenvalue sum) plotted against gradient number. The second or third gradient corresponds to the clearest elbow in the plot.

**Supplementary Figure 3:**
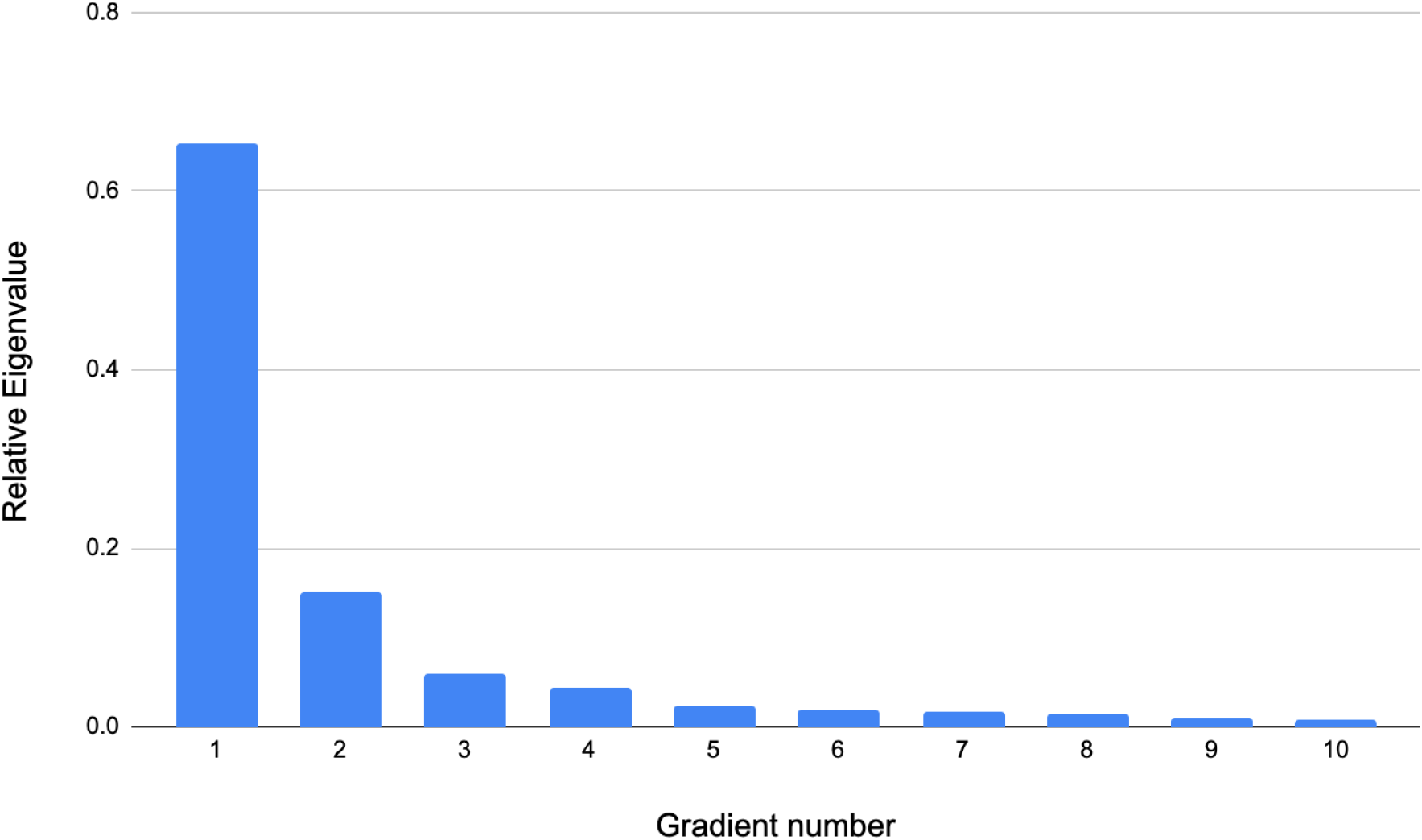
Scree Plot for Right Hippocampus FC gradients in Exploration Dataset. The second or third gradient corresponds to the clearest elbow in the plot.

**Supplementary Figure 4:**
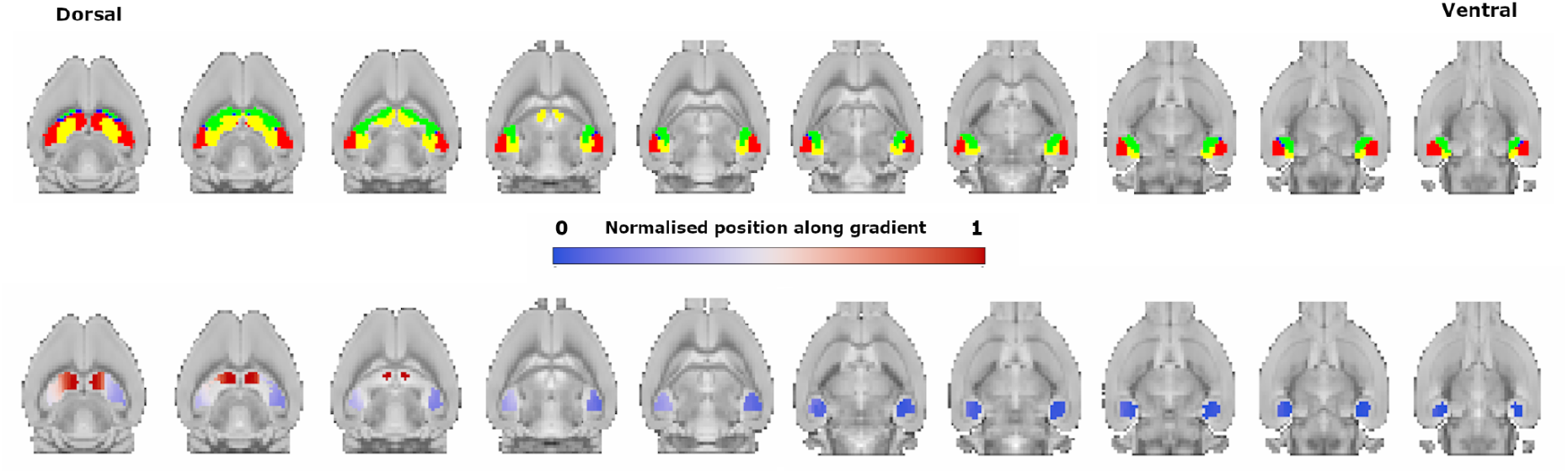
Principal FC Gradient in Validation Dataset. In the principal FC gradient in the validation dataset, the left gradient transitions more gradually from the dorsal hippocampus to the ventral compared to the right gradient and the gradient in the exploration dataset.

**Supplementary Figure 5:**
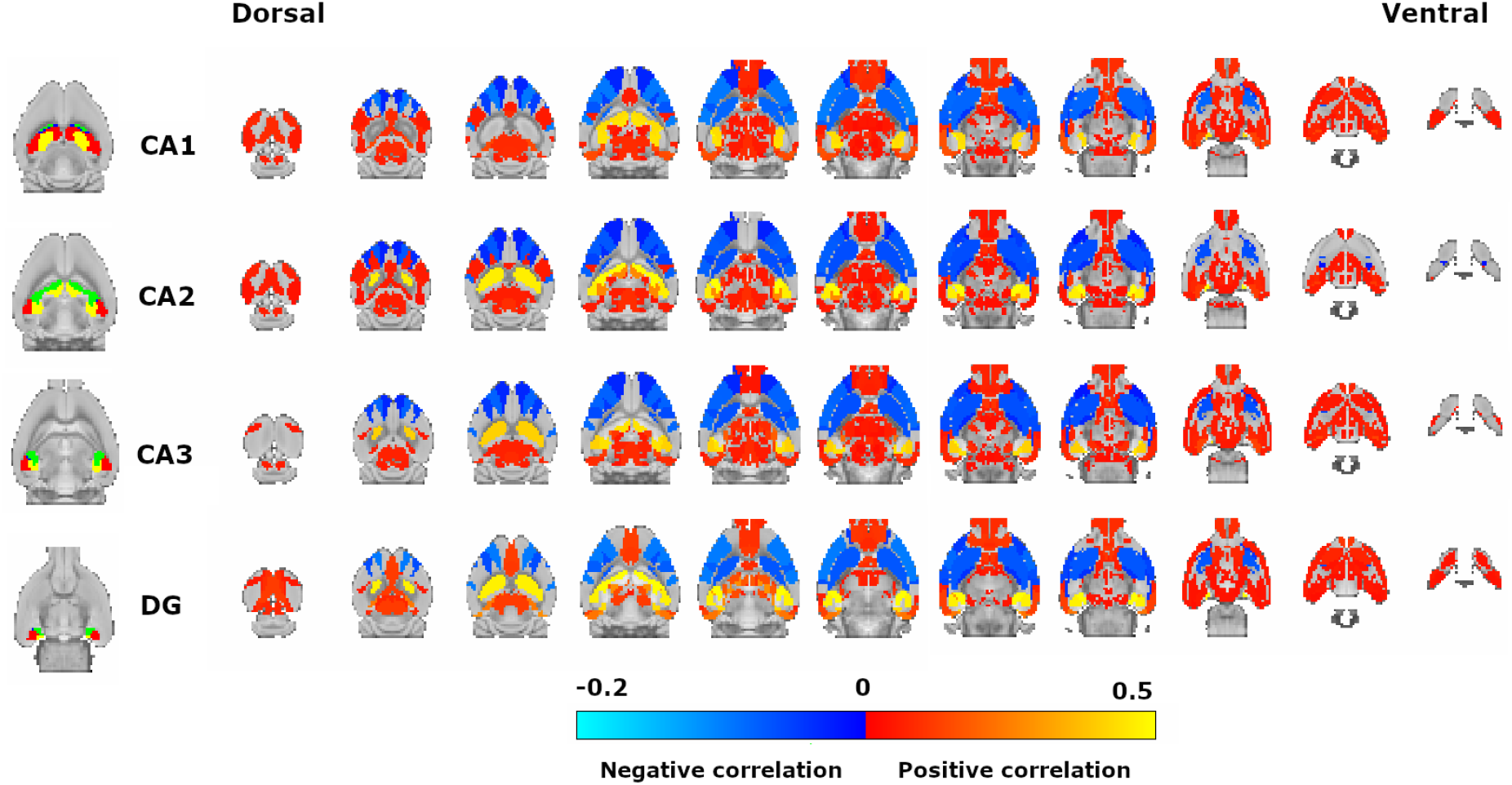
Significant Correlations of the Hippocampus Subfields. On the left, are the hippocampus subfields in the dorsal-to-ventral direction from top to bottom. The red region is the CA1 subfield, the blue region is the CA2 subfield, the green region is the CA3 subfield and the yellow region is the DG. Each row shows significant correlations (Pearson *R*) for a hippocampal subfield.

**Supplementary Figure 6:**
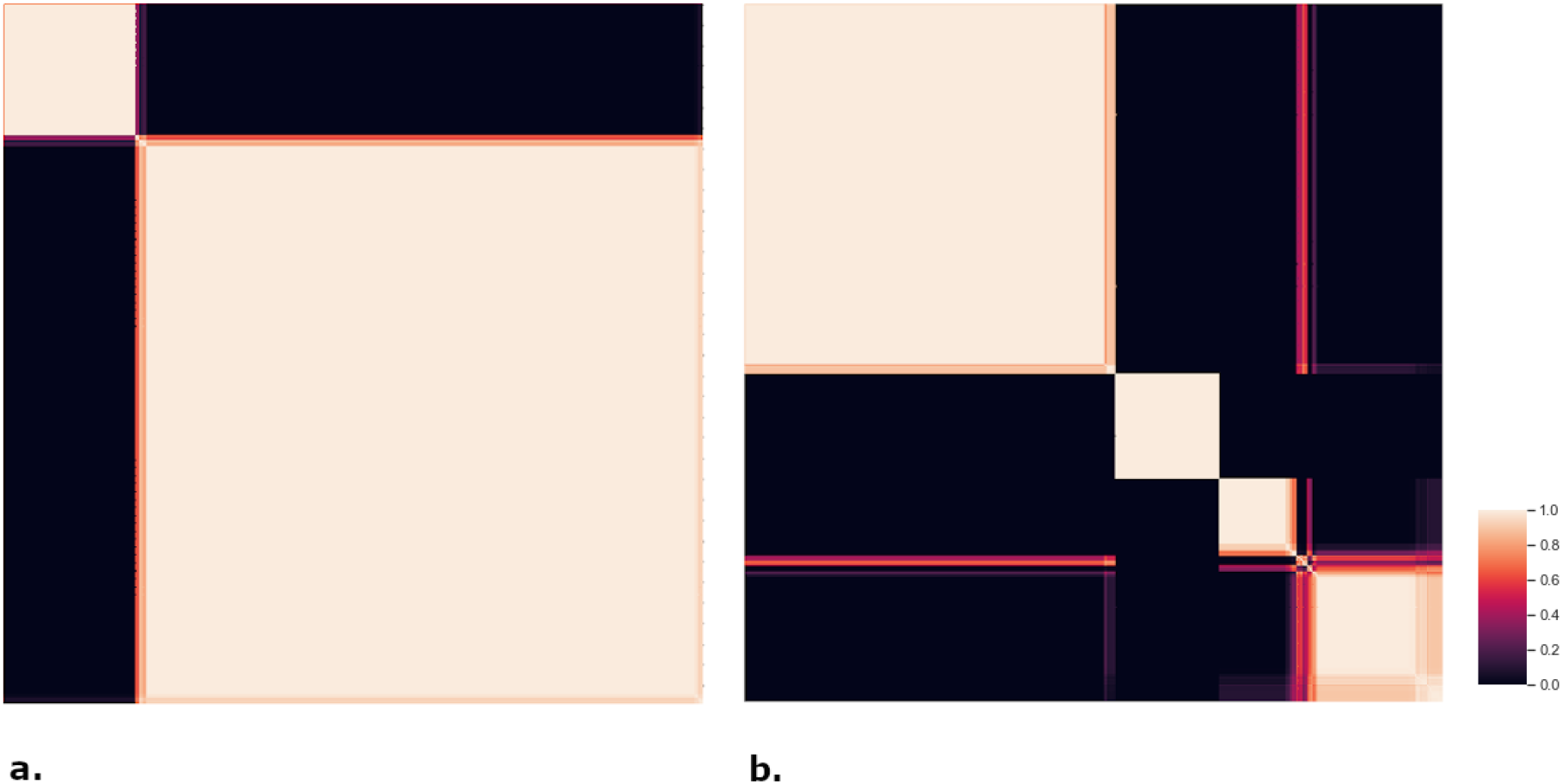
Heatmaps of Principal Gradient Clustering. Heatmaps for optimal clustering solutions for the principal gradients. Heatmaps are rearranged consensus matrices where voxels assigned to the same cluster are adjacent to each other. A value of 1 means a voxel pair is always assigned to the same cluster, 0 a voxel pair is never assigned to the same cluster. a. shows the heatmap for a two cluster solution for the left gradient, featuring relatively crisp boxes on the diagonal. b. shows the heatmap for a four cluster solution for the right gradient, featuring relatively crisp boxes on the diagonal.

**Supplementary Figure 7:**
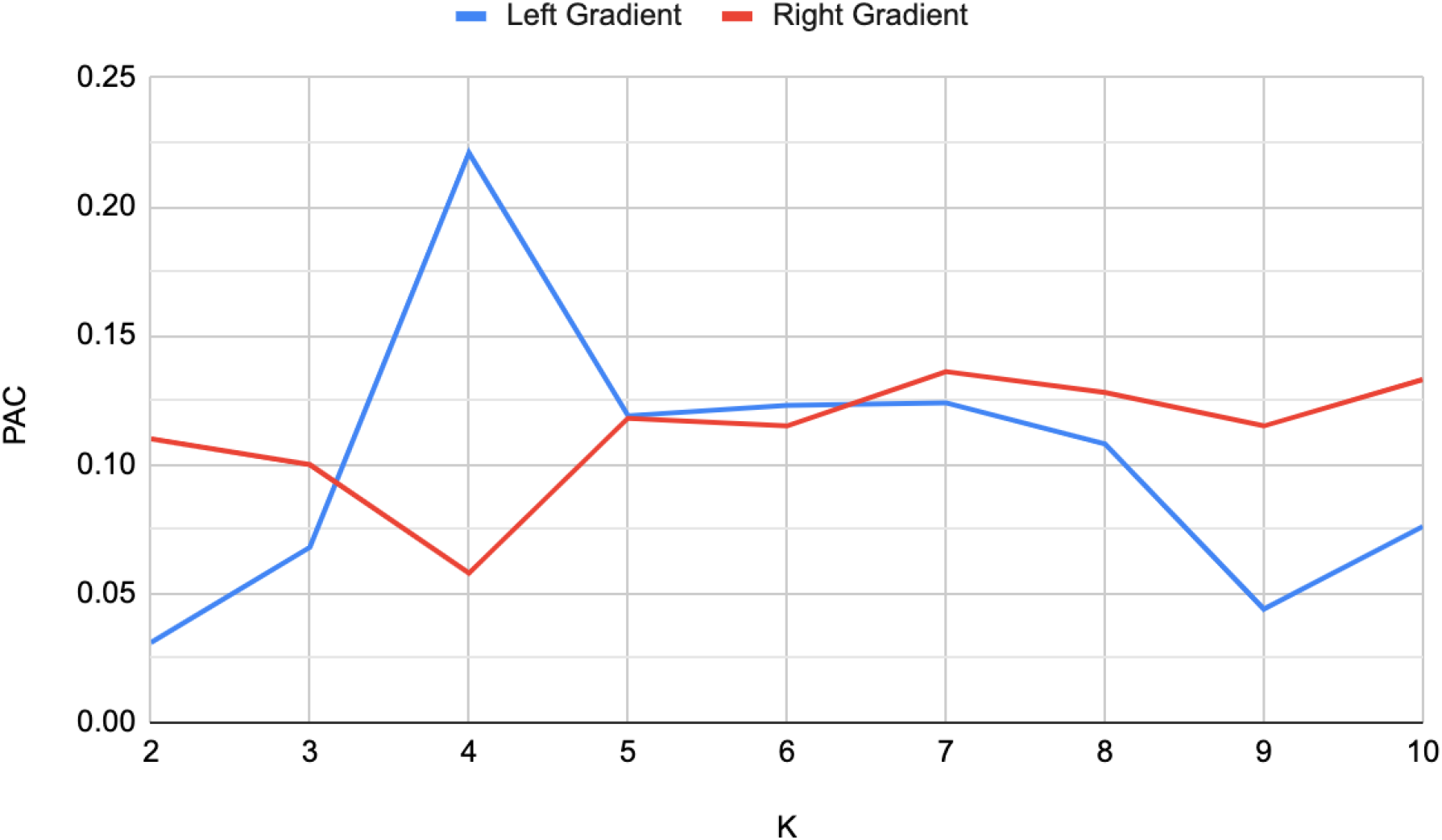
Principal FC Gradient PAC. PAC for the clustering solutions (K) of the left and right principal FC gradients.

**Supplementary Figure 8:**
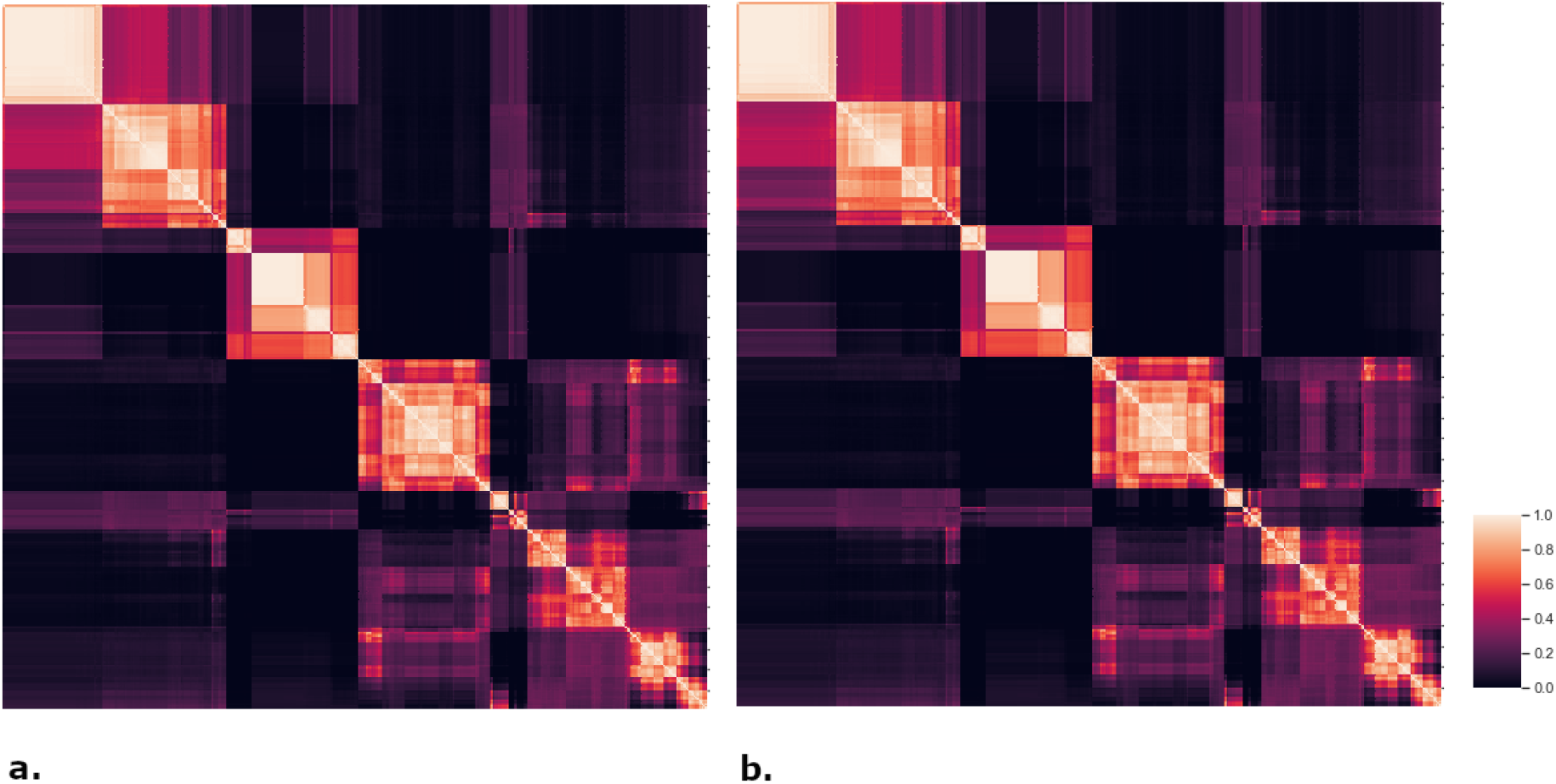
Heatmaps of Second Gradient Clustering. Heatmaps for a six clustering solution for the second gradient. a. shows the heatmap for a six cluster solution for the left gradient, featuring relatively uncrisp boxes on the diagonal. b. shows the heatmap for a six cluster solution for the right gradient, featuring relatively uncrisp boxes on the diagonal.

**Supplementary Figure 9:**
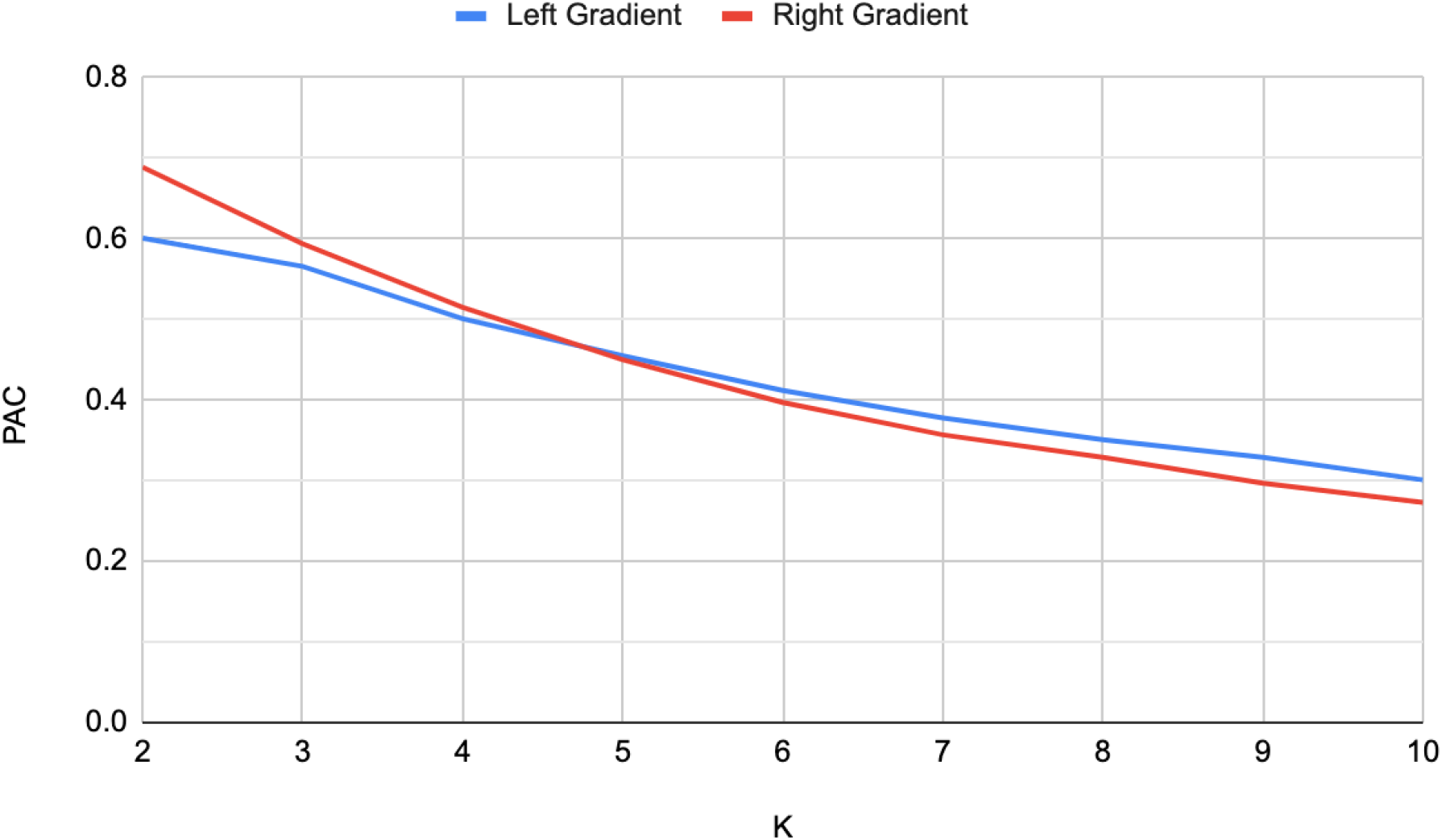
Second FC Gradient PAC. PAC for the clustering solutions (K) of the left and right second FC gradients.

**Supplementary Figure 10:**
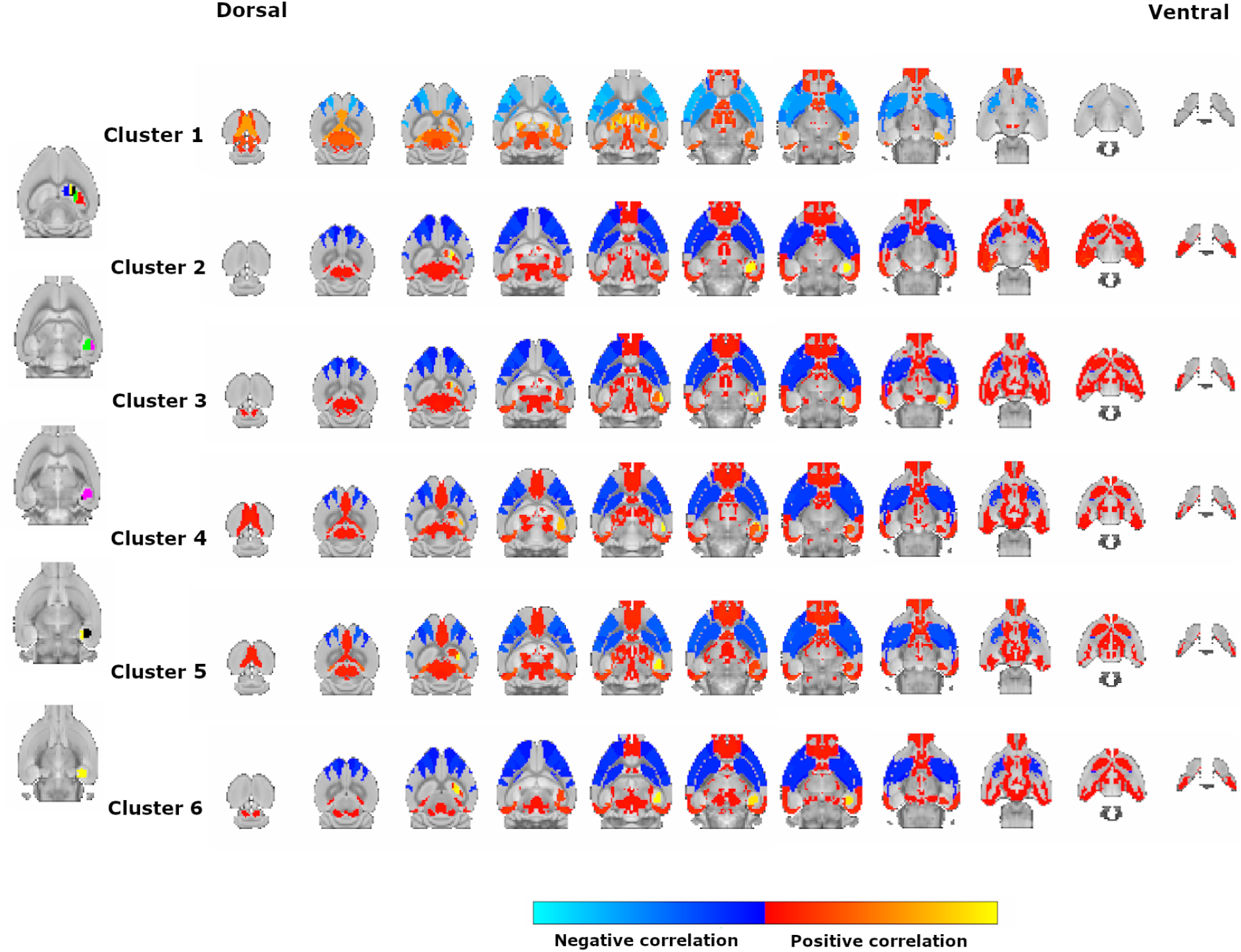
Significant Correlations of the Right Second FC Gradient. The figure shows the different patterns of correlations across the gradient clusters for the second FC gradient of the right hippocampus (cluster 1 - blue, cluster 2 - yellow, cluster 3 - black, cluster 4 - green, cluster 5 - red, cluster 6 - pink)

**Supplementary Figure 11:**
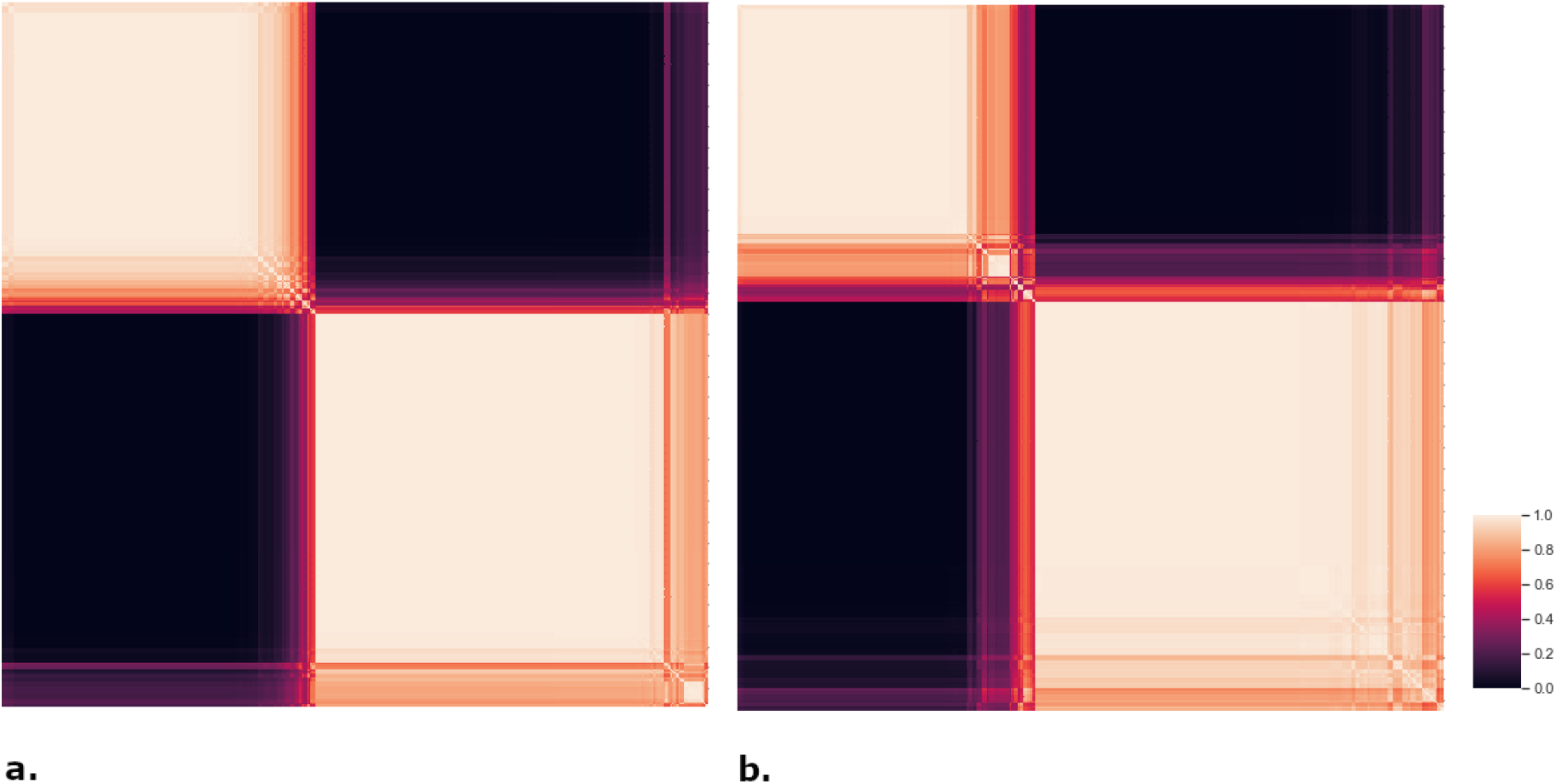
Heatmaps of the Third Gradient Clustering. Heatmaps for optimal clustering solution for the third gradient. a. shows the heatmap for a four cluster solution for the left gradient, featuring relatively crisp boxes on the diagonal. b. shows the heatmap for a four cluster solution for the right gradient, featuring relatively crisp boxes on the diagonal.

**Supplementary Figure 12:**
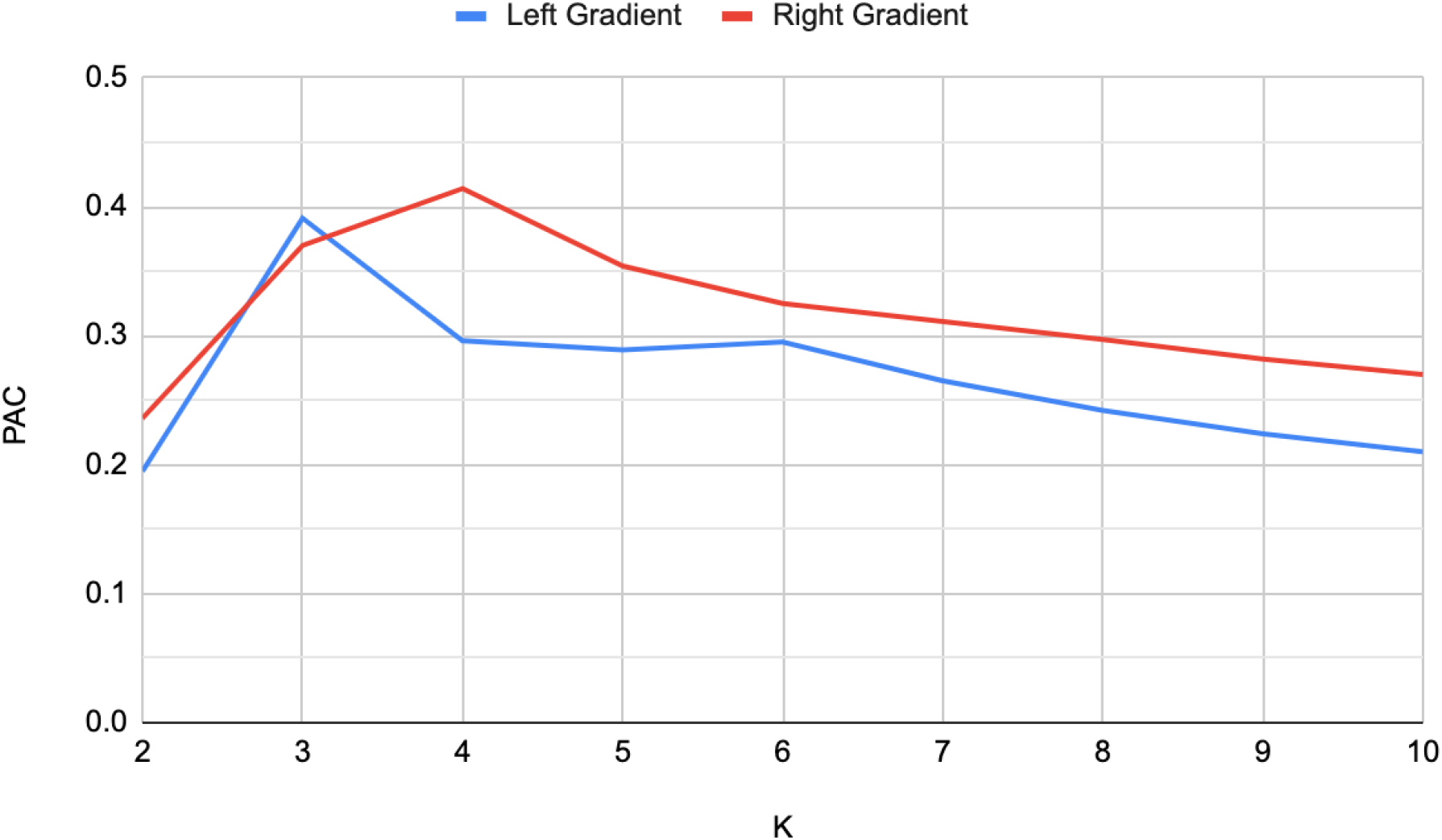
Third FC Gradient PAC. PAC for the clustering solutions (K) of the left and right third FC gradients.

**Supplementary Figure 13:**
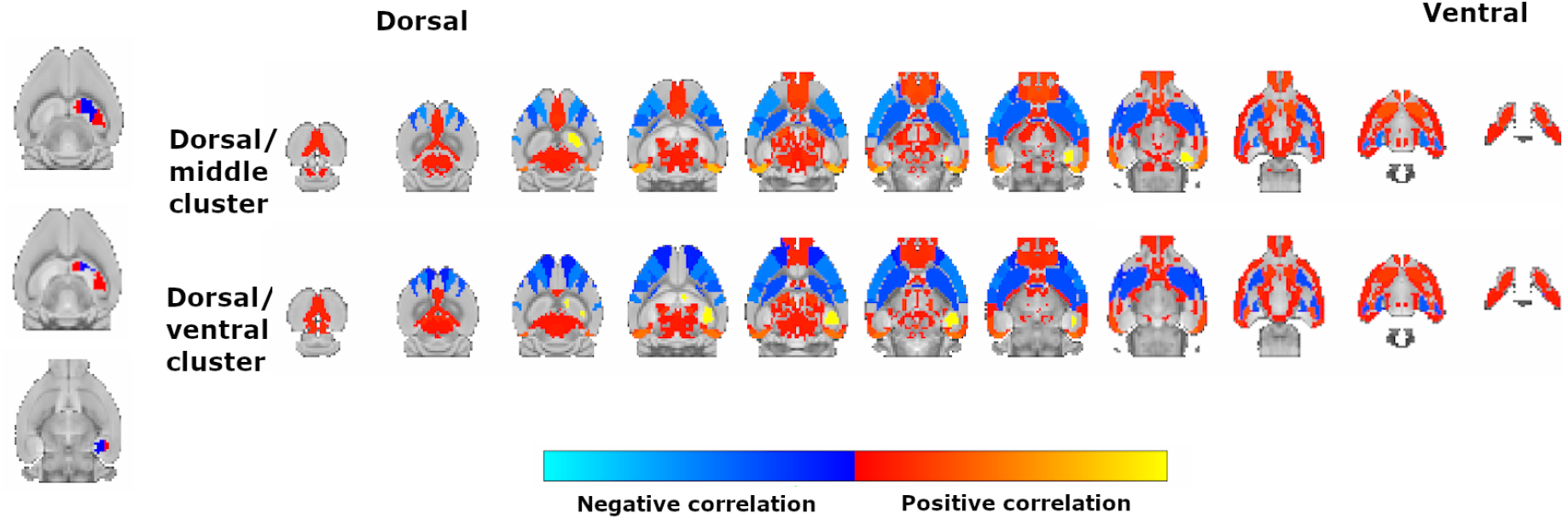
Significant Correlations of the Right Third FC Gradient. The figure shows the different patterns of correlations across the gradient clusters of the right third FC gradient.

**Supplementary Figure 14:**
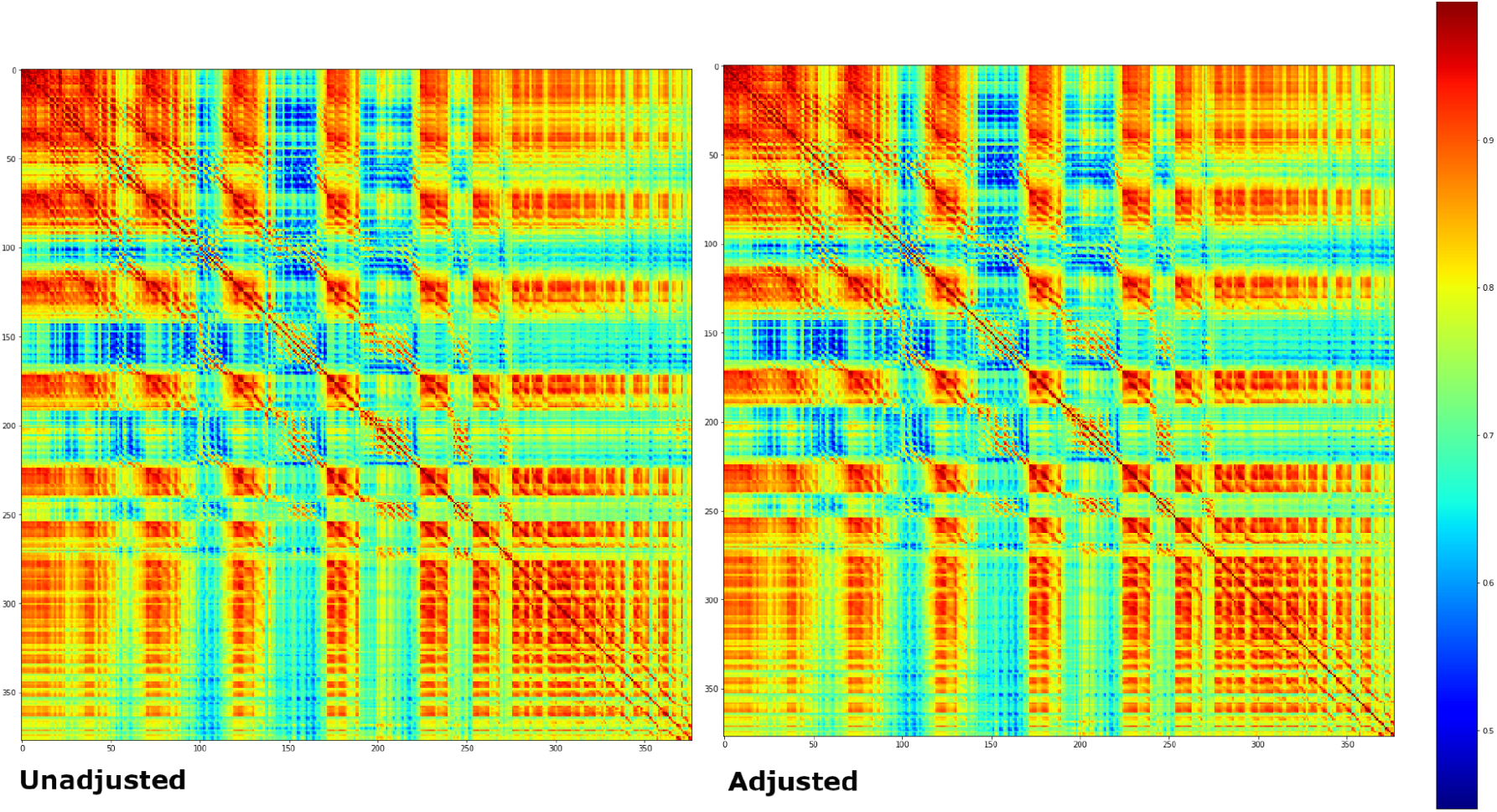
Unadjusted and Adjusted GE Similarity Matrices Left Hippocampus. Figure showing the unadjusted and adjusted gene expression similarity matrices for the left hippocampus. Elements are colour coded according to the colour bar on the right, where a warmer colour signifies a value closer to 1 and a cooler colour signifies a value closer to ∼0.45

**Supplementary Figure 15:**
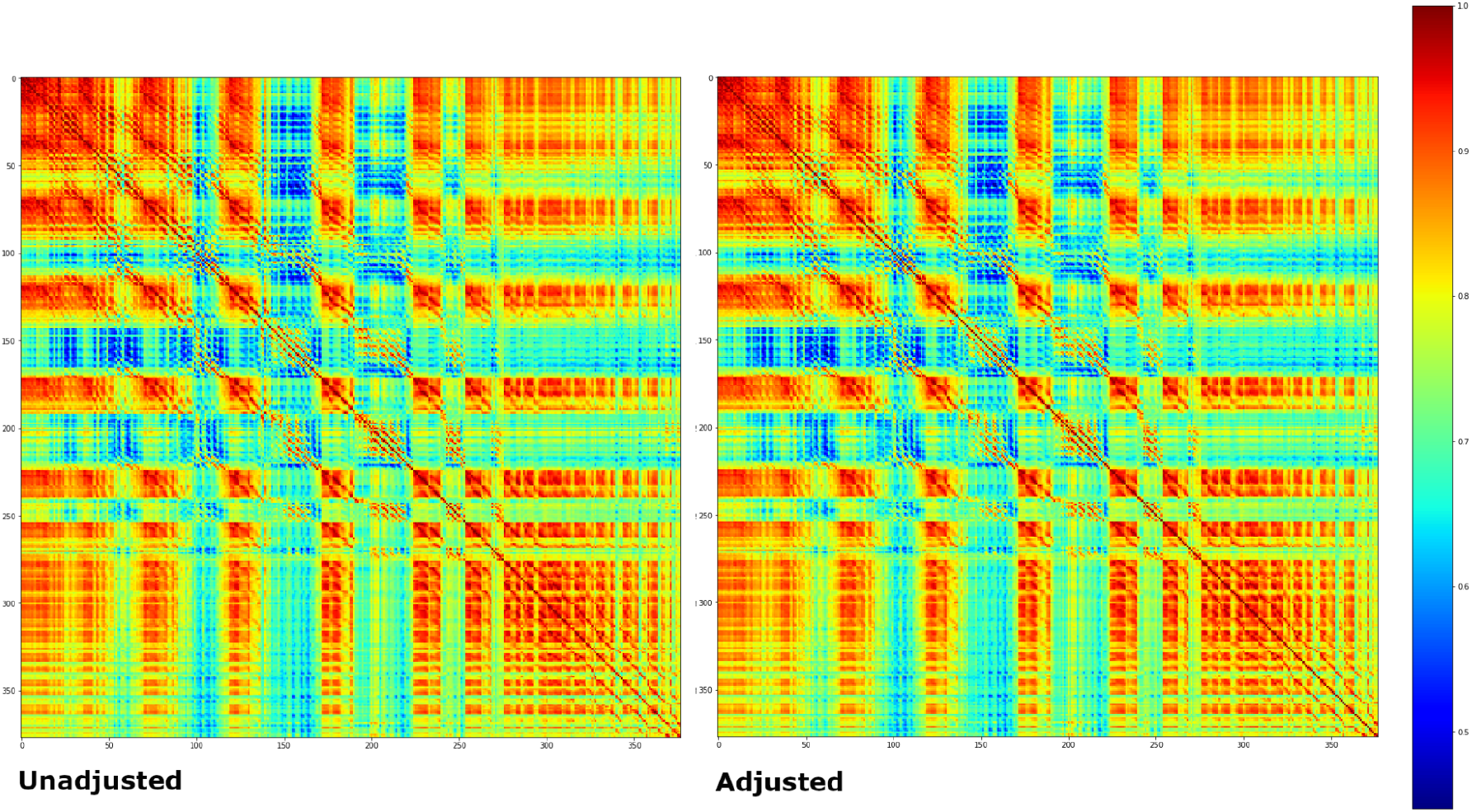
Unadjusted and Adjusted GE Similarity Matrices Right Hippocampus. Figure showing the Unadjusted and adjusted gene expression similarity matrices for the right hippocampus.

**Supplementary Figure 16:**
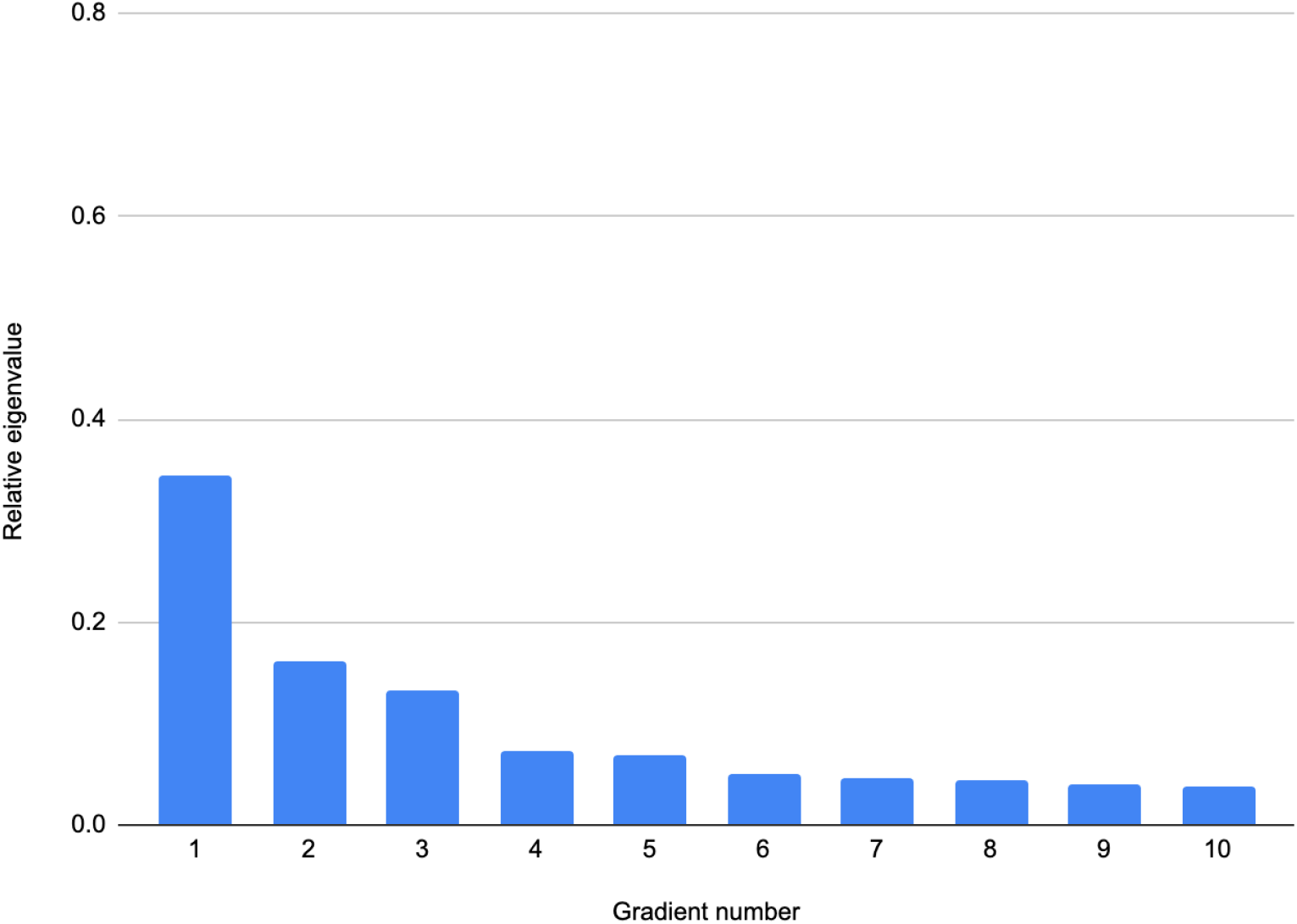
Scree Plot for Left Hippocampus GCE Gradients. Relative eigenvalue (reciprocal of eigenvalue divided by total eigenvalue sum) plotted against gradient number. The third gradient corresponds to the clearest elbow in the plot.

**Supplementary Figure 17:**
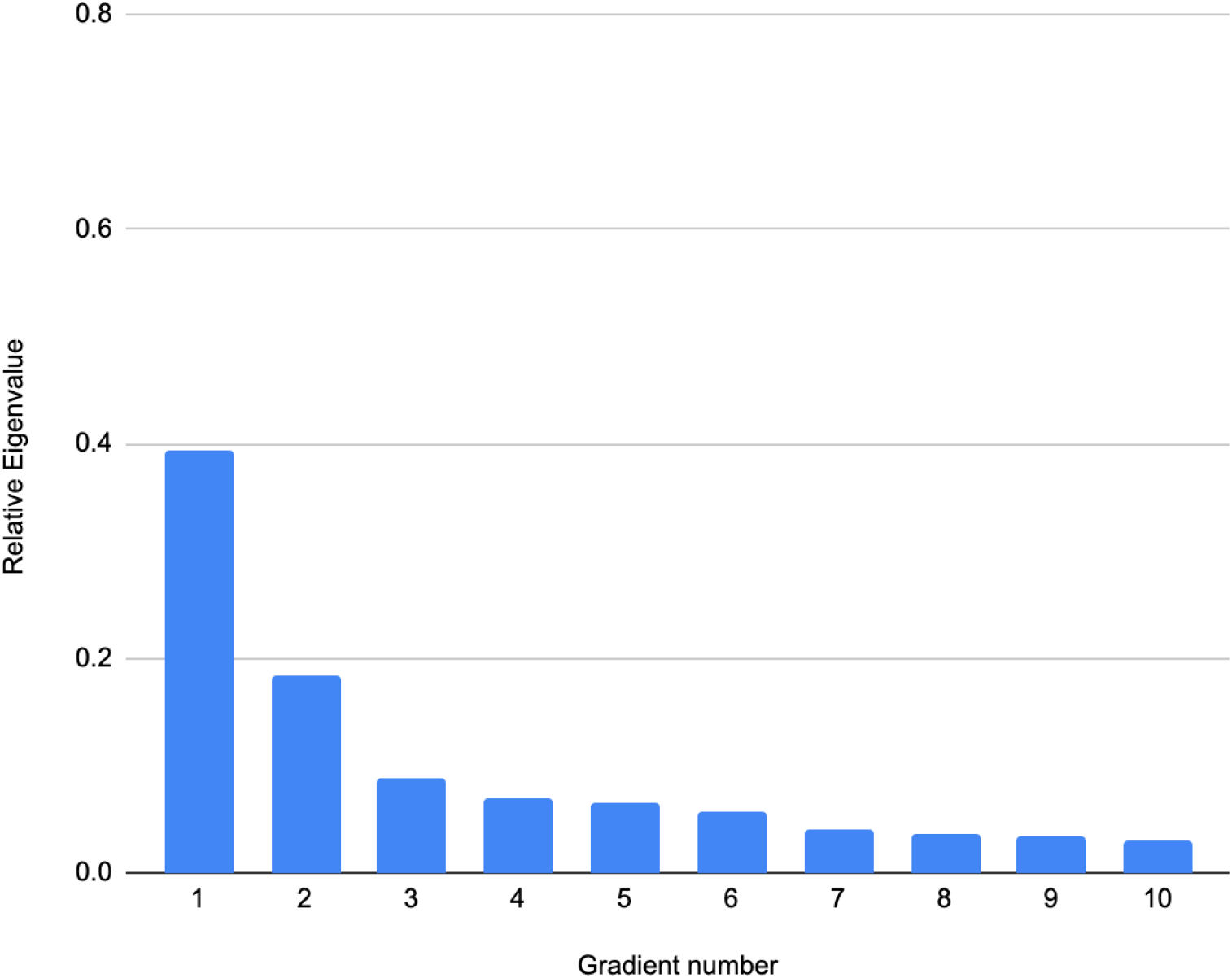
Scree Plot for Right Hippocampus GCE Gradients. The third gradient corresponds to the clearest elbow in the plot.

